# Altered MAM function shifts mitochondrial metabolism in SOD1-mutant models of ALS

**DOI:** 10.1101/2022.09.22.508778

**Authors:** Delfina Larrea, Kirstin A. Tamucci, Khushbu Kabra, Kevin R. Velasco, Taekyung D. Yun, Marta Pera, Jorge Montesinos, Rishi R. Agrawal, John W. Smerdon, Emily R. Lowry, Anna Stepanova, Belem Yoval-Sanchez, Alexander Galkin, Hynek Wichterle, Estela Area-Gomez

## Abstract

Mitochondrial defects are a common hallmark of familial and sporadic forms of amyotrophic lateral sclerosis (ALS). However, the origin of these defects, including reduced pyruvate metabolism and reduced oxygen consumption, is poorly understood. These metabolic functions are regulated in specialized endoplasmic reticulum (ER) domains in close contact with mitochondria, called mitochondrial-associated ER membranes (MAM). Recently it has been shown that MAM domains are disrupted in ALS, but the connection between MAM dysregulation and mitochondrial defects in ALS cells remains unclear. Using human embryonic stem cell (ESC)-derived motor neurons (hMNs) and mouse models with ALS-pathogenic mutations in superoxide dismutase 1 (SOD1), we found that the glycolytic deficiency in ALS is a direct consequence of the progressive disruption of MAM structure and function that hinders the use of glucose-derived pyruvate as a mitochondrial fuel and triggers a shift in mitochondrial substrates from pyruvate to fatty acids. This glycolytic deficiency, over time, induces significant alterations in mitochondrial electron flow and in the active/dormant (A/D) status of complex I in spinal cord, but not in brain. These data agree with a role for MAM in the maintenance and regulation of cellular glucose metabolism and suggest that MAM disruption in ALS could be the underlying cause of the bioenergetic deficits observed in the disease.

## Introduction

Amyotrophic lateral sclerosis (ALS) is a fatal neurological disorder characterized by the selective loss of motor neurons (MNs), resulting in muscle atrophy, paralysis, and respiratory failure (Rowland and Shneider, 2001). Most ALS cases have an unknown etiology, and are referred to as sporadic ALS (sALS); however, a growing number of mutations in more than 20 genes have been described and associated with familial ALS (fALS) (Marangi and Traynor, 2015). These include mutations in TAR DNA binding protein 43 (TDP43, gene *TARDBP*), guanine nucleotide exchange factor C9orf72 (gene *C9orf72)*, fused in sarcoma (FUS, gene *FUS*), and superoxide dismutase-1 (SOD1, gene *SOD1*), with SOD1 mutations accounting for almost 20% of fALS cases (Robberecht and Philips, 2013). The diverse functions of these genes implicate numerous cellular pathways and pathogenic processes in ALS progression. Yet, common to all forms of the disease, mitochondrial dysfunction has been shown to precede the loss of MNs and has been suggested to play a central role in the development of the disease, but the nature of this bioenergetic dysfunction is still unclear (Smith et al., 2019). Specifically, ALS mitochondria display significant alterations in their structure and dynamics, as well as defects in oxidative phosphorylation (OxPhos) and ATP production (Smith et al., 2019), primary due to a decreased activity of complex I (CI) of the respiratory chain (Ghiasi et al., 2012; Singh et al., 2021; Wiedemann et al., 2002).

Complex I is the main entry point of electrons into the respiratory chain for ATP production and is an essential hub for the regulation of metabolic signaling in response to changes in nutritional status (Hirst, 2013) and cellular redox state (Drose et al., 2014). Under physiological oxygen concentrations (normoxia), pyruvate derived from the breakdown of glucose is the most efficient mitochondrial fuel source for sustaining the high energetic demands of many cell types, including neurons (Belanger et al., 2011); cardiomyocytes are a notable exception to this rule, as cardiomyocyte mitochondria preferentially oxidize fatty acids (FAs) for ATP production (Stanley et al., 2005). Upon entry into mitochondria, pyruvate is converted to acetyl-CoA (by the pyruvate dehydrogenase complex, PDHC), which then enters the tricarboxylic acid (TCA) cycle for the subsequent production of reducing equivalents. NADH is oxidized by CI to NAD^+^ and the resulting electrons are transferred to complex III (CIII) by coenzyme Q (CoQ) followed by transport to complex IV (CIV) by cytochrome *c*. Electron transfer is coupled to the pumping of protons (H^+^) from the matrix to the intermembrane space by CI, CIII and CIV, allowing for the generation of the proton gradient that ultimately drives ATP synthesis by complex V (CV; ATP synthetase [ATPase]). Note that CII (also called succinate dehydrogenase [SDH]) activity also contributes to the function of the respiratory chain. At CII, two linked redox reactions take place: oxidation of succinate to fumarate, with the concomitant reduction of covalently-bound FAD to FADH_2_, and then oxidation of FADH_2_ to FAD, with the concomitant reduction of CoQ to CoQH_2_. However, succinate oxidation by CII does *not* result in the translocation of protons into the intermembrane space and therefore yields fewer ATP molecules compared to NADH oxidation (Nicholls, 2013; Speijer, 2011).

When the availability of glucose-derived pyruvate is limited, other substrates, such as fatty acids (FAs) and amino acids (AAs), are used as carbon sources for ATP production (Divakaruni et al., 2017; Hasselbalch et al., 1994; Mergenthaler et al., 2013; Owen et al., 1967; Worth et al., 2014). Compared to glucose, oxidation of FAs, and to a lesser extend AAs, results in increases in succinate and its oxidation to fumarate, as well as the in the reduction and oxidation of enzymes that contain covalently-linked FAD^+^ cofactors, such as CII, and other enzymes electron transferrin flavoproteins, Acyl-CoA dehydrogenase and glycerol-3-phosphate dehydrogenase. Oxidation of FAs increases the flow of electrons from CII to CoQ, bypassing CI (Nicholls, 2013; Speijer et al., 2014) and promoting respiration through CII. Consequently, the prevalence of FA oxidation over glucose decreases the ratio of NADH to FADH_2_ electron flux (Speijer et al., 2014) and, as explained above, yields fewer ATP molecules compared to pyruvate oxidation (Nicholls, 2013; Speijer, 2011).

Under normal nutritional conditions, electrons from both CI and CII arriving at CoQ are transferred forward to CIII. However, under nutritional overload (i.e. a high-fat diet), the excess of FADH_2_ over NADH exceeds the capacity of the reduced CoQ pool to transfer electrons to CIII. In this situation, the “excess” electrons take the path of least resistance and begin to flow backwards to CI, a process called reverse electron transfer (RET) (Nicholls, 2013). Notably, the oxidation of succinate can also induce elevations in the pool of reduced CoQ (i.e. CoQH_2_) and induce RET (Chance and Hollunger, 1960).

Under RET conditions, NAD^+^ is *reduced* by CI, boosting NADH pools (Robb et al., 2018) at the expense of a higher membrane potential. This process is associated with the highest rates of superoxide (O_2_^-^) production by CI, and the activation of superoxide dismutases, in order to mitigate the production of reactive oxygen species (ROS) (Pryde and Hirst, 2011; Robb et al., 2018). To diminish ROS generation under these conditions, CI undergoes a conformational transition from an active (A) to a dormant (D) state, thereby arresting RET, most probably to lessen ROS-mediated oxidative damage under conditions of RET (Drose et al., 2016). Furthermore, the build-up of NADH under RET conditions decreases the NAD^+^:NADH ratio that can inhibit the activity of glycolytic enzymes (the Randle cycle (Randle et al., 1963)) (Berthiaume et al., 2019) and the oxidation of FAs (Bartlett and Eaton, 2004).

Notably, defects in glucose metabolism are a common and early event across all forms of ALS (Pradat et al., 2010). Specifically, reductions in glucose uptake and in the expression of glucose transporters have been described numerous times in ALS patients and in brain and spinal cord (SPC) from animal models analyzed at both pre- and post-symptomatic stages (Tefera et al., 2021); increased glycogen deposits in postmortem spinal cord ALS sections (Dodge et al., 2013); reduction in insulin sensitivity (Reyes et al., 1984); and increased circulation of fatty acids in the serum of ALS patients (Pradat et al., 2010), although the source of these defects is unclear. Relevant to this reduction in glucose metabolism, recent work has revealed the contribution of a specific domain of the ER, called mitochondria-associated membranes (MAM), to the regulation of glucose and pyruvate metabolism (Theurey and Rieusset, 2017). MAM is a transient lipid raft domain formed by local and temporary increases in the levels of cholesterol in the ER, where multiple enzymatic activities converge to coordinately regulate several key pathways in response to metabolic changes (Vance, 2014). Importantly, impairments in the formation of MAM have been associated with the pathogenesis of several neurodegenerative diseases (Area-Gomez et al., 2012; Guardia-Laguarta et al., 2014; Larrea et al., 2019), including ALS (De Vos et al., 2012; Stoica et al., 2014; Watanabe et al., 2016).

In the present study, we aimed to elucidate the potential connection of MAM impairment to the alterations in glucose metabolism that occur in ALS. The results reported here indicate that, in the context of SOD1 mutations in animal and hESC-derived cell models of ALS, impaired regulation of MAM in neurons disrupts pyruvate metabolism and triggers a shift to FAs as an alternative mitochondrial fuel. Over time, this metabolic shift results in the induction of RET and the inactivation of CI. Overall, our data agree with a role for MAM in the maintenance and regulation of glucose metabolism and suggest that its disruption is the origin of the bioenergetic deficits observed in the disease.

## Results

### Mitochondrial respiratory defects in ALS^SOD1^ models are progressive and substrate-dependent

Mitochondrial bioenergetic deficiency is a common phenotype in all forms of ALS, but its cause is still unclear (Crugnola et al., 2010; Smith et al., 2019). Notably, multiple reports have implicated SOD1 as a mediator of the mitochondrial disturbances in the disease, both when mutated in fALS, or misfolded in the context of sALS (Allen et al., 2014; Bartolome et al., 2013; Kawamata et al., 2017).

To understand the origin of these metabolic alterations, we first confirmed that the transgenic (Tg) mouse model of ALS used here (overexpressing human *SOD1* carrying the G93A, mutation [SOD1^G93A^]) indeed displayed mitochondrial bioenergetic deficiencies in the brain and spinal cord (SPC) compared to age-matched non-transgenic (NTg) controls. We performed double histochemical staining to visualize the activities of complex IV (CIV, cytochrome *c* oxidase) and complex II (CII, succinate dehydrogenase), a method that enables the detection of cells with mitochondrial dysfunction *in situ* (Bonilla et al., 1992; Franco-Iborra and Tanji, 2020). In agreement with previous data obtained in ALS muscle (Scaricamazza et al., 2020), we found that brain and SPC samples from SOD1^G93A^ animals presented with defects in the respiratory chain at a pre-symptomatic stage (P60) (**Fig. 1**).

**Fig. 1.**
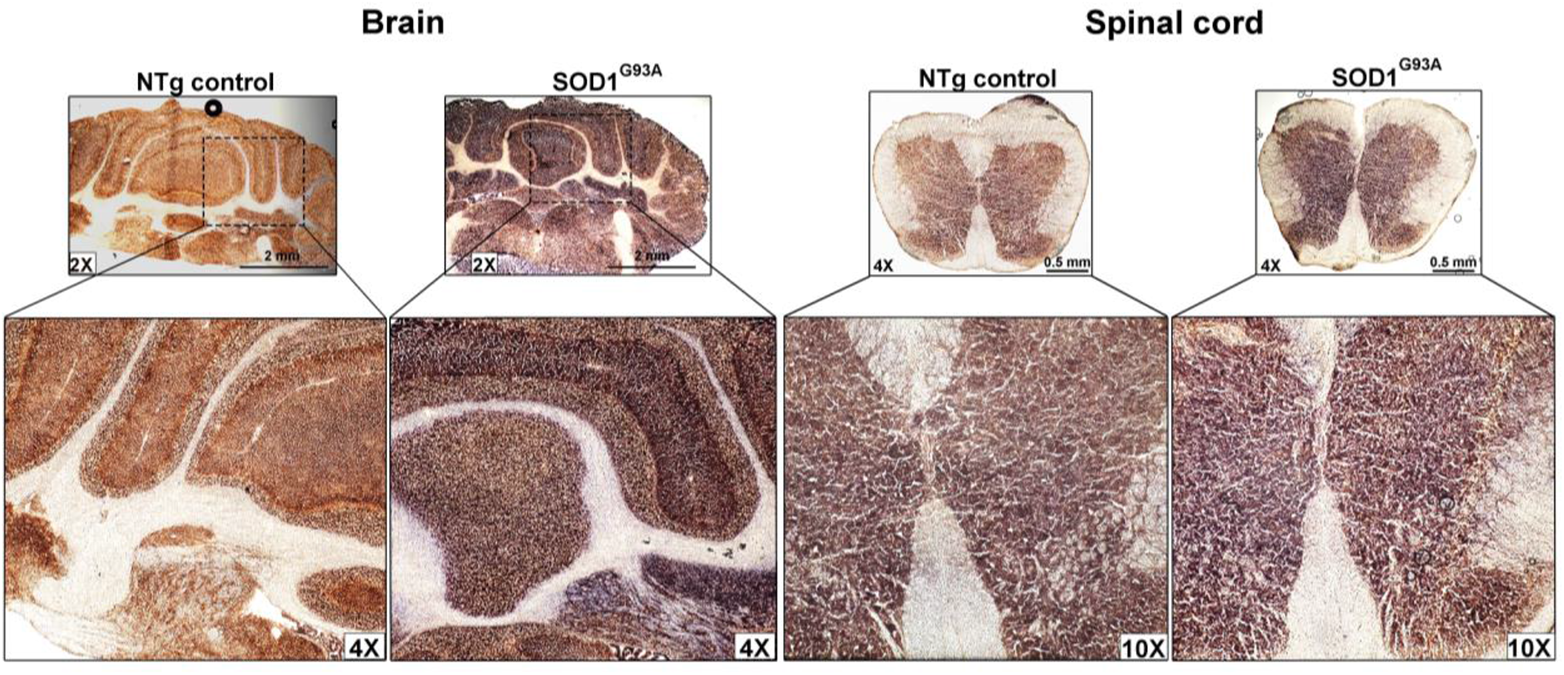
Defects in mitochondrial respiratory complexes in tissues from ALS^SOD1^ model mice. Cytochrome *c* oxidase (COX) and succinate dehydrogenase (SDH) activities are shown by double staining in 10-m-thick coronal sections of brain at the level of the cerebellum area and in cross sections of SPC tissues from SOD1^G93A^ and NTg control mice at pre-symptomatic stage P60. Note that, in control sections, the purple stain representing SDH activity is hidden by the brown stain representing COX activity, whereas the purple stain is prominently visible in mutant tissues. This suggests reduced COX activity in mutant samples that enables visualization of the SDH stain. The images shown are representative of 3 independent animals per group.

To further probe into the nature of these bioenergetic alteration(s), we measured oxygen consumption rates (OCR) in mitochondria isolated from brain and SPC of SOD1^G93A^ mice and NTg controls. To avoid the confounding effects of significant neuronal death, and of astro-/microgliosis, on oxygen consumption that occurs after P60 (Hall et al., 1998), we performed our analyses on mitochondria isolated from tissues at different pre-symptomatic ages (P15, P30, P60) and compared the results to analyses conducted at disease onset (P90) and late (P120) stages (**Figs. 2A, 2D and S1A, and supplementary Table 1**). We first measured respiration in isolated mitochondria using pyruvate (in the presence of malate to block CII activity) as a CI-supporting substrate whose oxidation activates NADH-complex I (CI)-driven oxygen consumption (NADH-OCR) and produces higher levels of NADH compared to the oxidation of other substrates, such as FAs and AAs (Schonfeld and Reiser, 2013; Speijer, 2011) (**Figs. 2A and 2B**). We then measured respiration using succinate as the substrate (in the presence of rotenone to inhibit CI activity), whose oxidation produces higher levels of FADH_2_ (i.e., reduces the covalently-bound FAD+ cofactor in FAD-containing enzymes) compared to the oxidation of pyruvate (Schonfeld and Reiser, 2013), and thus, activates FADH_2_-complex II (CII)-driven oxygen consumption (FADH_2_-OCR) (Martinez-Reyes and Chandel, 2020) (**Figs. 2C and 2D**).

**Fig. 2.**
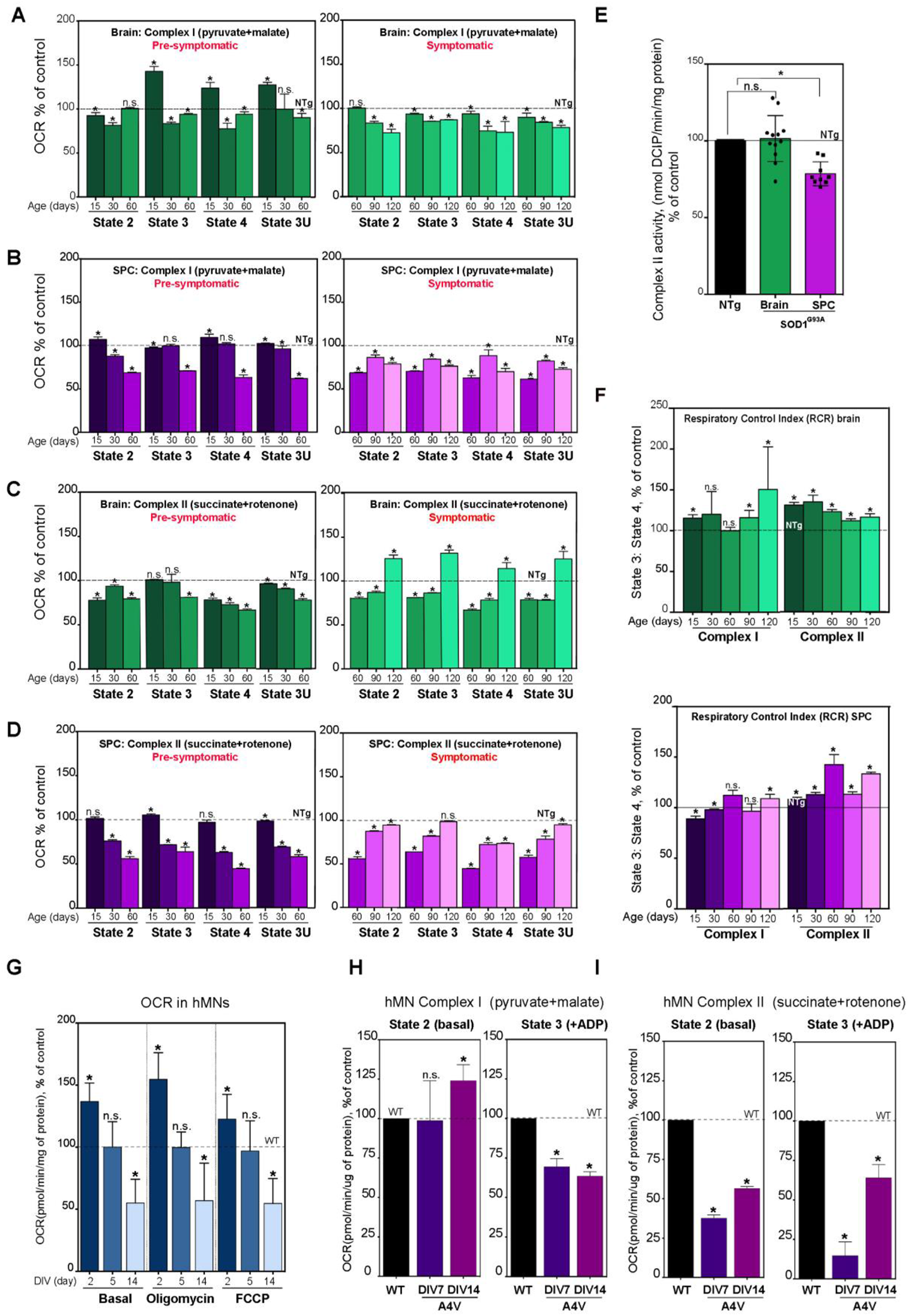
Progressive alterations in mitochondrial respiration in SOD1^G93A^ mice. (**A-D**) Quantification of the oxygen consumption rate (OCR, in pmole O_2_/min/mg protein) in mitochondria isolated from the brain (**A** and **C**) and spinal cord (SPC) (**B** and **D**) from SOD1^G93A^ mice analyzed at the indicated ages (in days) compared to aged-matched NTg controls (set at 100%; dotted lines). In each assay 6 µg of mitochondria and 8 µg of mitochondria were loaded per well to measure CI and CII, respectively. Respiration was measured at baseline or **State 2** (i.e. initial respiration in the presence of CI or CII added substrates (i.e. pyruvate+malate or succinate+rotenone, respectively); after the addition of ADP to initiate **State 3**; after the addition of oligomycin to initiate **State 4**; and after the addition of the uncoupler, FCCP, to initiate **State 3-uncoupled (3U)**; Note that CI-driven oxygen consumption (NADH-OCR) declines after P60 (**A** and **B**) and CII-driven respiration (FADH_2_-OCR) increases after P60 (**C** and **D**). (n=3 biological replicates using 3 - 5 technical replicates in each experiment. Data are shown as the average ± SD; *p <0.05, t-test). (**E**) Complex II enzyme activity in isolated mitochondria from brain and SPC from mutant animals at P60 relative to the NTg control average (set at 100%; dotted line). (Average ± SD of n=3-5 biological replicates using 3-4 technical replicates in each experiment; *p <0.05, t-test). (**F**) Respiratory control ratio (RCR; ratio of state 3:state 4) calculated from the respiratory values from Fig. 2 A-D, in brain **(top panel)** and SPC **(bottom panel**) from SOD1^G93A^ mice relative to the NTg control average (set at 100%; dotted lines). Note that in both brain and SPC, the RCR due to FADH_2_-OCR was increased above control at all ages analyzed, contrary to what was observed for NADH-OCR. (**G**) OCR in human motor neurons carrying the A4V mutation (hMNs^A4V^) relative to that in WT controls (set at 100%; dotted lines) in the presence of pyruvate as a mitochondrial substrate. Note that basal OCR decreased after day 5 of differentiation *in vitro* (DIV) compared to WT, as it did after addition of oligomycin (Oligo) and FCCP. Data are shown as the average of n=3-5 biological replicates using 5 technical replicates. Average ± SD; *p <0.05, t-test). (**H-I**) NADH- and FADH_2_-OCR in permeabilized hMNs^A4V^ relative to the WT control average (set at 100%; dotted lines). Notation as above. (Average of n = 3-5 technical replicates ± SD; *, p<0.05, t-test).

At the earliest stage examined (P15), mitochondria isolated from SOD1-mutant brain showed a slight reduction in NADH-OCR at baseline (state 2), but significantly increased OCR after stimulation with ADP (state 3), and a sustained elevation in additional assayed states (states 4 and 3U) (**Fig. 2A**). This elevation was not observed at later pre-symptomatic stages (P30 and P60), nor at symptomatic stages (P90 and P120) (**Fig. 2A**). Compared to brain, and as expected for a motor neuron disease, NADH-OCR of SPC mitochondria showed a progressive decline in in OCR at pre-symptomatic stages that was sustained after symptoms appeared (**Fig. 2B**). We note that the declines in NADH-OCR were in the range of 30-50%, as seen previously by others (Mattiazzi et al., 2002; Wiedemann et al., 2002).

We then measured FADH_2_-OCR and observed a dramatically different trend in both brain and SPC, especially at later stages of the disease. In brain samples, similar to NADH-OCR, there was a progressive reduction in FADH_2_-OCR (30-50%) that was maintained at disease onset (P90), but increased dramatically over NTg levels during the end-stages (P120) (**Fig. 2C**). In SPC, the progressive decline in FADH_2_-OCR began earlier (at P30) and was more exacerbated (50-60%) than was the decline in NADH-OCR at P60 (**Fig. 2D**). Also, as in brain, we observed that FADH_2_-OCR increased up to NTg levels during the end-stages, (**Fig. 2D**), perhaps due to a significant proliferation of glial cells in mutant tissues. Thus, mutant brain tissues, presented with similar slight reductions in NADH- and FADH_2_-OCR at pre symptomatic and early onset stages, below a pathogenic threshold (~25% less than NTg control). Unexpectedly, SPC presented with more dramatic and earlier reductions in FADH_2_-OCR compared to NADH-OCR in SPC, brain tissues and NTg controls.

To determine whether the defects in FADH_2_-OCR in SOD1-mutant cells were associated with defects in the intrinsic CII enzymatic activity, we measured succinate dehydrogenase enzymatic activity in brain and SPC mitochondria at a presymptomatic (P60) stage. Despite the reductions in FADH_2_-OCR observed in SOD1^G93A^ tissues (**Figs. 2A and 2D**), CII activity was not significantly altered in brain and only mildly reduced in SPC (**Fig. 2E**). Moreover, the changes in OCR did not correlate with changes in the levels of mitochondrial respiratory chain components (**Fig. S1B**), with alterations in mtDNA content (**Fig. S1C**), or with changes in mitochondrial biogenesis markers (**Fig. S1D**). This result implies that, despite being enzymatically intact, the decline in CII-driven oxygen consumption in the SOD1^G93A^ mice suffers a more pronounced decline than does CI-driven OCR, even before disease onset. This is a surprising conclusion, given that the central nervous system has historically been deemed to rely almost exclusively on NADH (i.e. CI)-linked substrates (Mergenthaler et al., 2013); and thus, OCR defects are expected to have a more significant impact on CI as compared to CII.

To further examine these mitochondrial phenotypes, we calculated the respiratory control ratio (RCR) (i.e. state 3:state 4) under both substrates (pyruvate and succinate) as a measure of the coupling efficiency between mitochondrial respiration and ATP production (Brand and Nicholls, 2011). RCR values showed only slight differences between controls and mutant samples when pyruvate was used as the substrate (NADH-OCR). However, succinate oxidation (FADH_2_-OCR) resulted in significantly higher RCR in mitochondria isolated from both SOD1^G93A^ brain and SPC (**Fig. 2F**). A higher RCR implies a greater capacity for ATP production (Scialo et al., 2017), which suggests that brain and SPC mitochondria from SOD1 mice use succinate in a slightly more efficient manner than they do pyruvate or other NADH-linked substrates.

Since motor neurons (MNs) are the most affected cell type in ALS, we next assessed the mitochondrial respiratory profile (in the presence of pyruvate and glucose) of non-permeabilized cultured human embryonic stem cell (ESC)-derived motor neurons (MNs) carrying a mutation in SOD1 (SOD1^+^/^A4V^; denoted as hMNs^A4V^) and WT controls at different days *in vitro* (DIVs) after differentiation (**Fig. 2G**). This approach evaluates OCR in intact cells, mimicking respiratory conditions in the cell and providing information on cellular respiratory control (i.e. ATP production rate, proton leak rate, coupling efficiency, maximum respiratory rate, RCR and spare respiratory capacity) (Brand and Nicholls, 2011). Mutant hMNs^A4V^ initially displayed elevated OCRs at DIV2 that declined over time to ~50% of WT hMNs at DIV14 (**Figs. 2G and S1E**) as shown by others (Ludtmann et al., 2017; Szelechowski et al., 2018). In addition, mutant hMNs had a lower spare respiratory capacity (SRC; uncoupled respiration minus basal respiration) compared to controls, indicative of an impairment in mitochondrial metabolic flexibility (**Fig. S1F**). Contrary to what was found in the SOD1-mutant brain and SPC, the defects in hMN mitochondrial respiration were associated with an increase in mitochondrial biogenesis (as measured by mtDNA content and *PPARCG-1A* mRNA levels, **Fig. S1G**) and an increase in mitochondrial mass (as measured by TOM20 protein levels, **Fig. S1H**).

To allow for the entry of exogenous ADP and specific oxidizable substrates (pyruvate and succinate) to feed electrons into respiratory chain, we repeated our respirometry analysis in permeabilized hMNs. As before, mutant hMNs^A4V^ at both DIV7 and DIV14 fed with pyruvate (NADH-OCR) showed essentially no alteration in OCR at baseline, but progressive reductions in state 3 OCR (after ADP addition) (**Fig. 2H**). Notably, the reduction in NADH-OCR was only evident after the addition of ADP (state 3) when respiration is driven by ATP hydrolysis, but not with non-phosphorylating respiration (state 2, no ADP) (Nicholls, 2013). The surprising finding that SOD1-mutant cells are unable to respond effectively to pyruvate-driven ADP stimulation, while still maintaining normal OCR at state 2, is consistent with a scenario in which electrons derived from NADH substrates are not used efficiently for ATP production and trigger a leakage of protons, a mechanism that mitigates ROS production and mitochondrial damage at the expense of ATP production (Nicholls, 2013; Tabassum, 2020). On the other hand, and similar to SOD1^G93A^ tissues, respiratory defects were more exacerbated when succinate was used as the substrate, and these defects appeared even at baseline before the addition of ADP (**Fig. 2I**).

The difference between the respiration values using pyruvate vs. succinate points to a fundamental alteration in the flow of electrons in ALS mitochondria. In particular, these metabolic changes suggest that the flux via CII to oxygen is greater in ALS than in normal cells. This could in turn, indicate a shift from NADH-linked substrates towards FADH_2_-linked fuels, such as fatty acids (FAs), for the production of ATP in the context of SOD1 mutations.

### A shift to fatty acid consumption induces reverse electron transfer and the transition of CI to the dormant state in SOD1^G93A^ mice

Defects in the metabolism of glucose (i.e., the use of pyruvate) and increased oxidation of fat sources have been observed frequently in ALS patient samples, cells, and mouse models (Palamiuc et al., 2015; Steyn et al., 2020; Tefera et al., 2021). In support of these findings, our results indicate that hexokinase (HK) activity, the first step in glycolysis, was reduced progressively in brain and SPC from SOD1^G93A^ mice compared to controls (**Fig. 3A**). Similarly, we found significant reductions in the activity of PDHC, which converts pyruvate into acetyl-CoA to feed the TCA cycle, in both mutant tissues, even at pre-symptomatic stages (**Fig. 3B**). Likewise, hMNs^A4V^ presented with gradual reductions in HK and PDHC activities compared to WT controls (**Fig. 3C**), confirming the known decline in glucose metabolism in ALS (Tefera et al., 2021). In all cases, mRNA expression for these genes was essentially unchanged compared to controls (Fig. S2).

**Fig. 3.**
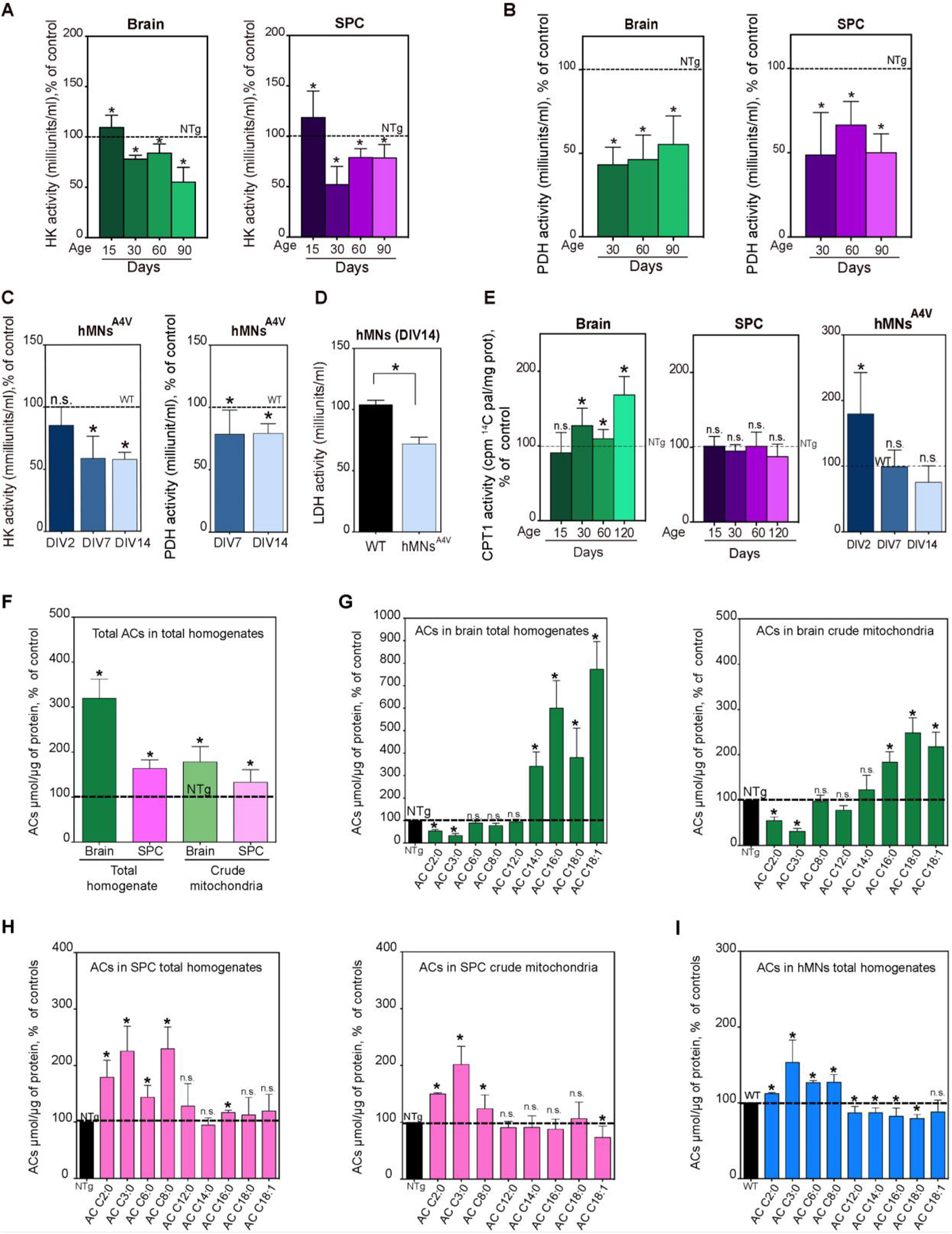
Defects in glycolysis and pyruvate metabolism in ALS models shifts mitochondria towards fatty acid oxidation. The activities of **(A)** hexokinase (HK) in total homogenates and **(B)** pyruvate dehydrogenase (PDH) in crude mitochondrial fractions were decreased in brain and SPC from SOD1^G93A^ mice at the indicated ages, and in **(C)** hMNsA4V after the indicated DIV compared to WT controls (set at 100%; dotted line). **(D)** Lactate dehydrogenase (LDH) activity was reduced in hMNs^A4V^ vs WT. **(E)** CPT1 activity was increased in brain mitochondria over time, unchanged in SPC, and decreased in hMNs^A4V^ compared to WT controls. **(F-I)** Lipidomics analysis in brain and SPC mitochondria from SOD1^G93A^ mice showed significant increases in the levels of total acylcarnitines (ACs) compared to NTg controls **(F)**, due mainly to increased ACs containing C_14_-C_18_ in brain **(G)**, while conversely, the AC species that were elevated in SPC **(H)** and hMNs^A4V^ **(I)** were shorter and enriched in C_2_-C_8_. All of the above suggest a reduction in the production and oxidation of pyruvate, in agreement with the reduction in glucose metabolism, and a shift towards fatty acid utilization. All assays were averages of 3-5 biological replicates each consisting of 3-5 technical replicates. Data are shown as mean ± SD. Statistical comparisons are made to NTg (for mouse experiments) or WT (for hMN experiments) controls (two-tailed t-test; α=0.05; *, p<0.05).

Neuronal pyruvate pools are derived mainly from astrocytic lactate. Points of regulation in this process include neuronal uptake of astrocyte-secreted lactate and the conversation of lactate to pyruvate by lactate dehydrogenase (LDH) (Belanger et al., 2011). We therefore measured LDH activity in our hMN cultures and found significant decreases in the hMN^A4V^ mutants (**Fig. 3D**), despite showing higher expression levels of *LDHA* mRNA (**Fig. S2B**). Taken together, these results support a decline in the production and use of pyruvate in neural tissues of SOD1^G93A^ mice and hMNs^A4V^ compared to controls, and thus, a decrease in the utilization of pyruvate as a substrate for mitochondrial respiration catabolic degradation. In light of these data, we tested the use of fatty acids as an alternative mitochondrial fuel source in our models by measuring the activity of the mitochondrial carnitine palmitoyl-transferase 1 (CPT1). This enzyme conjugates carnitine to long-chain FAs (C_12_ to C_20_) for their translocation into mitochondria and subsequent oxidation (McGarry and Foster, 1974). We found that CPT1 activity was elevated in mitochondria isolated from brain SOD1^G93A^ mice, even at presymptomatic stages (**Fig. 3E**). Remarkably, however, SPC showed no significant changes in CPT1 activity compared to controls (**Fig. 3E**). On the other hand, hMNs^A4V^ showed a progressive reduction in CPT1 activity, even though *Cpt1* expression was unchanged (**Fig. S2C**).

To further examine this phenotype, we quantified the levels of FAs bound to carnitine (i.e., acyl-carnitines, ACs), by performing a lipidomics analysis on homogenates and mitochondrial fractions from SOD1^G93A^ tissues and controls. Both brain and SPC at P60 showed significantly higher total AC levels compared to controls (**Fig. 3F**). However, only brain showed specific elevations in the longer FA chains (AC_14_ to AC_18_) characteristic of increased usage of FAs as mitochondrial substrates and consistent with increased CPT1 activity (Adeva-Andany et al., 2019), while the concentrations of shorter AC species were reduced (**Fig. 3G**). Mirroring these results, SPC samples and hMNs^A4V^ displayed significantly increased levels of short ACs (AC_2_ to AC_8_), and no changes or even reductions in longer AC species (**Figs. 3H and 3I**). Notably, increases in short-chain ACs have been shown to arise from incomplete β-oxidation of FAs in mitochondria (Koves et al., 2008; Schooneman et al., 2013), which suggest that FA oxidation in mitochondria from mutant SPC was less efficient compared to brain.

As mentioned above, under conditions of increased FA oxidation, the levels of succinate and FADH_2_ increase alongside with a reduction of the covalently bound FAD^+^ cofactor of CII. A portion of CII-bound FADH_2_ electrons is then transferred back to CI to propel NAD^+^ reduction to form NADH (Acin-Perez et al., 2014; Guaras et al., 2016), a mechanism known as reverse electron transfer (RET) (Scialo et al., 2017). In healthy cells, RET is believed to be activated to maintain NADH levels, membrane potential, and ATP production under conditions of low pyruvate and/or excess succinate (Scialo et al., 2017). Under RET, cells display reduced NAD^+^:NADH ratios and increased membrane potential (Nicholls, 2013). Consistent with the activation of RET in the context of SOD1 mutations, our data revealed that mitochondria from SOD1^G93A^ brain and SPC at P60, as well as those from hMNs^A4V^, had significantly lower NAD^+^:NADH ratios compared to controls (**Fig. 4A**).

**Fig. 4.**
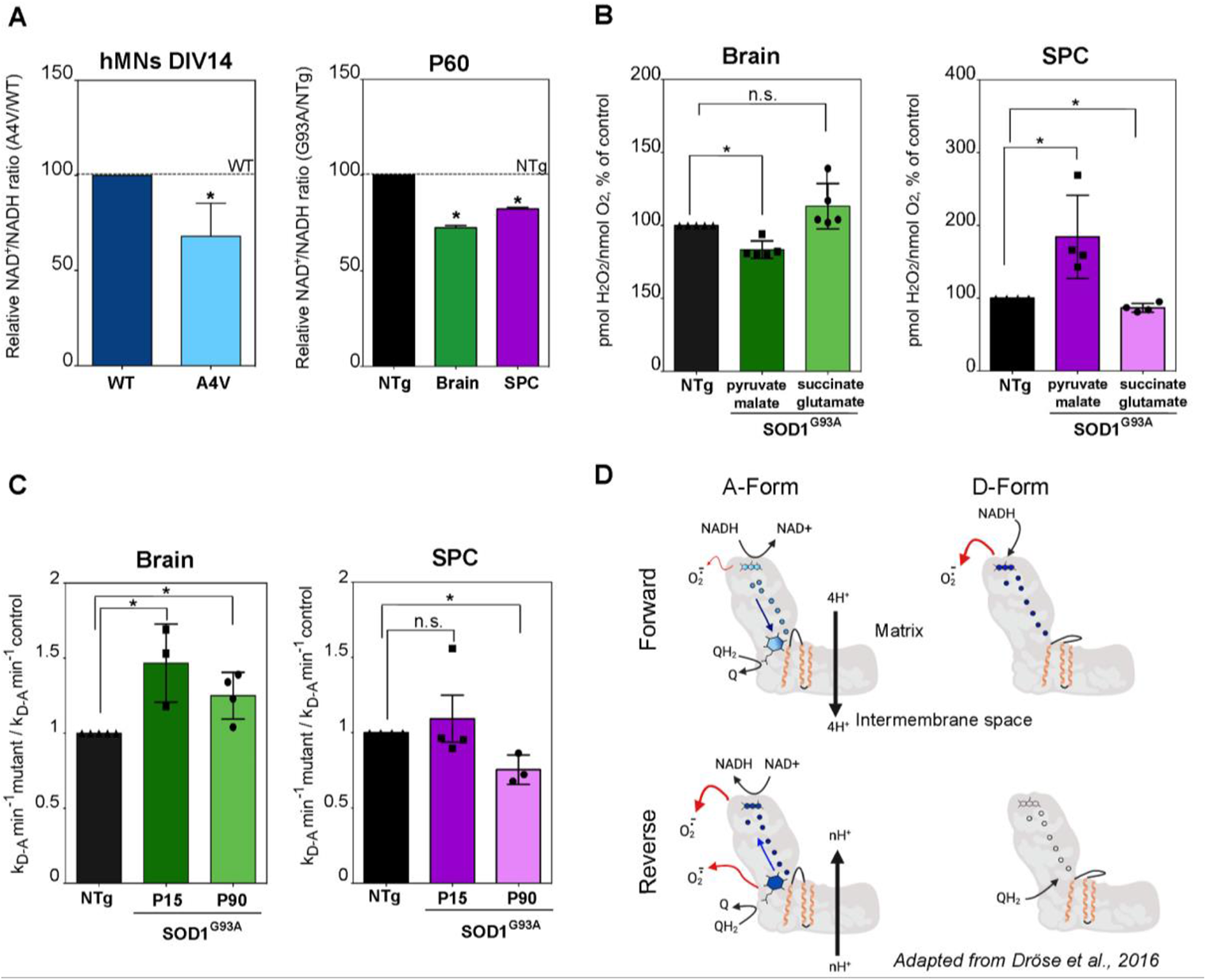
Defects in NADH levels and complex I activation in mitochondria from SOD1^G93A^ mice. **(A)** NAD^+^:NADH ratio in hMNs^A4V^ at DIV14 and in SPCs at P60 were reduced relative to controls (dotted line, set at 100%). (Average ± SD of n=3-4 biological replicates using 2 technical replicates each; two-tailed t-test; α=0.05; *, p<0.05). **(B)** Quantification of hydrogen peroxide produced during the oxidation of different substrates in mitochondria from SOD1^G93A^ mice vs NTg (set at 100%). (Average ± SD of 3-5 biological replicates using 3-4 technical replicates each; two-tailed t-test; α=0.05; *, p<0.05). **(C)** Analysis of the D-to-A transition constant for CI, calculated and expressed as min-1 (see **Fig. S3** for absolute values). Results are represented as values in SOD1^G93A^ mice vs NTg (set at 100%). (Average ± SD of 3-4 biological replicates using 3 technical replicates each; two-tailed t-test; α=0.05; *, p<0.05). **(D)** Model. Superoxide production in isolated mitochondria from SOD1^G93A^ animals can be attributed to the conformation of CI. CI can work in the forward mode, oxidizing NADH and pumping H^+^ out of the matrix and reducing CoQ, or in the reverse mode, reducing NADH and oxidizing CoQ. Under the latter scenario, H^+^ are allowed to return to the matrix from the intermembrane space to maintain mitochondrial membrane potential. The reverse mode induces ROS and can affect CI structure (likely at ND3 [Drose 2016] [orange subunit]), shifting CI from the A-form to the D-form. In our model CI from SOD1^G93A^ SPC display a k_D-A_ transition that is slower compared to that in controls (see **4C**), which suggests that CI is predominantly present in the D-form. This results in the generation of high ROS levels in the presence of pyruvate (see **Fig. 4B**). In agreement with a predominant CI-D conformation *in vivo* no ROS is produced in the presence of succinate, as explained in Dröse et al., 2016. Instead, SOD1^G93A^ brain CI display a k_D-A_ transition that is faster compared to controls, which suggests that CI is predominantly present in the A-form, resulting in the generation of low ROS levels similar to those in controls. FMN, FeS clusters and ubiquinone are shown schematically. The intensity of the blue color filling of the redox centers of CI reflects the degree of their reduction. This figure was created with BioRender.

The reduction of NAD^+^ to NADH that occurs during RET is also associated with high rates of ROS production (Robb et al., 2018; Stepanova and Galkin, 2020; Stepanova et al., 2019b). Therefore, if persistent, CI working in reverse mode results in increased ROS levels over time, which in turn is associated with significant reductions in CI activity (Acin-Perez et al., 2014; Guaras et al., 2016). Surprisingly, mutant mitochondria from SPC, but not brain, underwent a substantial increase in the production of H_2_O_2_ when pyruvate, but not succinate, was used as substrate (**Fig. 4B**). These data suggest that, contrary to what occurs in SPC, even under RET conditions and reduced NAD^+^:NADH ratios, ALS-brain mitochondria are able to maintain a pool of active CI in forward mode (i.e., transferring electrons from NADH to CoQ), generating low levels of ROS associated with the oxidation of NADH-linked substrates (Drose et al., 2016).

Of note, increased RET and subsequent increases in ROS can be attenuated by transient deactivation of CI via conformational changes (Drose et al., 2014). Hence, to corroborate our data, we assessed the kinetics of transition from the catalytically deactivated, or dormant, “D-form” of CI to the fully active “A-form” (Drose et al., 2014) in mitochondrial fragments from brain and SPC of the SOD1^G93A^ mice (**Fig. 4C**). Intriguingly, at both P15 and P90, brain mitochondria from the mutant mice showed significant increases in the rate of CI activation, whereas SPC mitochondria showed a significant decrease in CI activation, but only upon disease onset (P90) (**Figs. 4C and S3**). As with the data on ROS production, these data also imply that under RET conditions, ALS brain maintains or even augments the pool of active CI, whereas SPC mitochondria maintain a pool of predominantly dormant CI.

In summary, our data indicate that, in the context of SOD1 mutations, defects in the use of NADH substrates induce a shift towards the oxidation of FAs to generate ATP. This metabolic change results in elevations in FADH_2_-OCR (CII)-driven respiration and subsequent activation of RET. This stimulation of RET induces increases in ROS production and the deactivation of CI in SPC, but not in brain. Our data thus support previously observed reductions in CI activity in an ALS model (Panov et al., 2011), but do not support the idea that CI is somehow defective. Rather, we conclude that CI is fundamentally normal in ALS SPC, but is more abundant in the dormant form.

### Alterations in glucose metabolism in SOD1 models are induced by MAM disruption

The regulation of glucose metabolism and insulin signaling has recently been shown to be modulated by the formation of MAM domains in the ER that are in close apposition to mitochondria (Theurey and Rieusset, 2017). Defects in the formation of MAM result in disturbances in glucose metabolism and/or changes in the use of pyruvate as a mitochondrial substrate (Rieusset, 2018; Tubbs et al., 2018b). (Interestingly, a growing number of reports have suggested that ER-mitochondria crosstalk and the enzymatic activities localized at MAM are disrupted in the context of ALS (De Vos et al., 2012; Sakai et al., 2021; Stoica et al., 2014; Watanabe et al., 2016). Therefore, we asked whether the impairments in pyruvate metabolism found in our mutant models were the consequence of defects in the formation and/or activation of MAM domains.

We first measured ER-mitochondria crosstalk by quantifying the synthesis and transport of phospholipids between the two organelles, a well-established measure of MAM functionality (Montesinos et al., 2020; Vance, 1990). In this *dynamic* assay, we found no significant changes in MAM activity in the brain of SOD1^G93A^ mice compared to controls at all ages analyzed (**Fig. S4**). On the other hand, SPC samples from these mice showed a progressive decline in MAM activity compared to controls, reaching statistical significance at disease onset (P90) (**Fig. 5A**). Using the same approach, we also quantified MAM activity in our cultured hMN^A4V^ models and found significant reductions in MAM activity in hMNs^A4V^ cells compared to WT controls (**Fig. 5B**).

**Fig. 5.**
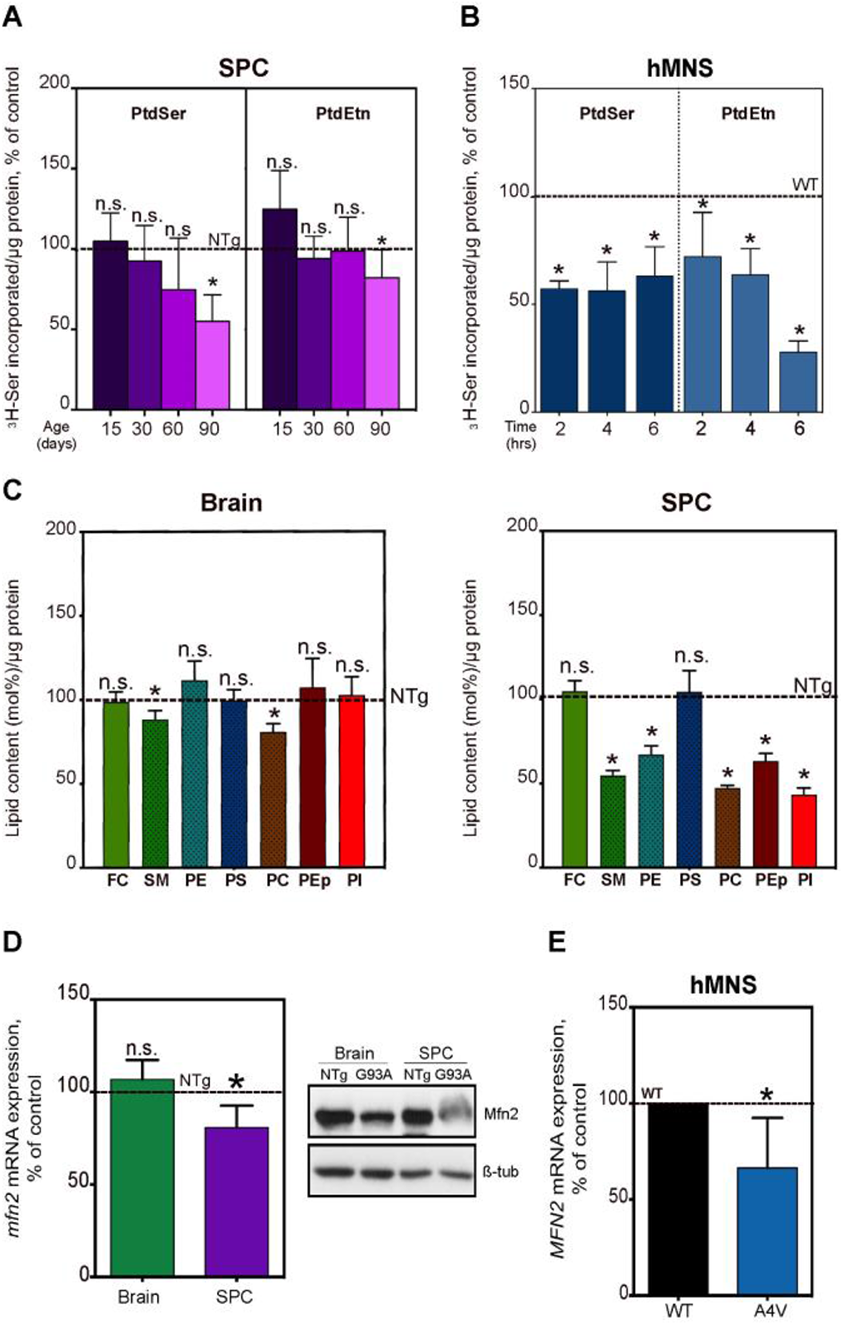
Downregulation of MAM activity in murine SOD1^G93A^ mice and hMNs^A4V^. **(A-B)** Quantification of MAM activity by the synthesis and transference of phospholipids between ER and mitochondria in SPC crude mitochondrial fractions from SOD1^G93A^ mice at the indicated ages (**A**) and in hMNs^A4V^ (DIV14) (**B**), relative to controls (dotted lines, set at 100%). (Average ± SD of n = 3-4 biological replicates; two-tailed T-test; α=0.05; *, p<0.05). **(C)** Lipidomics analysis of MAM fractions isolated from SOD1^G93A^ brain and SPC at pre-symptomatic stages (P60). FC: free cholesterol, SM: sphingomyelin, PE: phosphatidylethanolamine, PS: phosphatidylserine, PC: phosphatidylcholine, PEP: phosphatidylethanolamine plasmalogen, PI: phosphatidylinositol. (Average± SD values of n = 3 biological replicates using 3 technical replicates each; p* <0.05, t-test. (**D-E**) Quantification of mitofusin-2 (*Mfn2/MFN2*) gene expression by qRT-PCR and protein expression by WB in SOD1^G93A^ mice (**D**) and in hMNs^A4V^ at DIV14 **(E)**. (Average± SD values of n = 3 biological replicates using 3 technical replicates each; two-tailed t-test; α=0.05; *, p<0.05).

Similar to classic lipid raft membranes domains, MAM is a transient detergent-resistant domain that is enriched in cholesterol, sphingomyelin (SM) and saturated phospholipids, and whose formation and regulation depends on its lipid composition (Lingwood and Simons, 2010). Alterations in the lipid composition of MAM domains result in defects in the enzymatic activities localized and regulated at these ER regions (Area-Gomez et al., 2012; Pera et al., 2017). Thus, to validate MAM alterations in the context of mutations in SOD1, we assessed the lipid composition of MAM isolated from mutant and control brain and SPC by lipidomics (**Fig. 5C**). This analysis revealed a dramatic reduction in the steady-state levels of sphingomyelin (SM), phosphatidylcholine (PC), and phosphatidylethanolamine (PE) lipid classes in MAM fractions isolated from SPC, whereas in brain PE levels were unaffected and SM and PC were reduced only marginally (**Fig. 5C**). Our results indicate that a pathogenic mutation in ALS disrupts MAM function in SPC and MNs far more than it does in brain. In agreement with this idea, SPC (but not brain) from SOD1^G93A^ mice showed significant reductions in the expression of mitofusin 2 (gene: *Mfn2*), a MAM-localized protein that acts as an ER-mitochondria tether (de Brito and Scorrano, 2008; Naon et al., 2016), compared to controls (**Fig. 5D**). Likewise, the expression of *MFN2* was markedly reduced in hMNs^A4V^ compared to WT hMNs (**Fig. 5E**).

To relate these observations to the aforementioned alterations in glucose and pyruvate metabolism observed in ALS samples, we exploited the lipid-raft nature of MAM domains by inducing their formation in cultured hMNs. Specifically, MAM formation in the ER is triggered by local elevations in cholesterol delivered from the plasma membrane (PM) (Area-Gomez et al., 2012). Cholesterol mobilization has long been known to be stimulated by the activation of cellular sphingomyelinases (SMases) and can be replicated by incubation with exogenously added SMase (Slotte and Bierman, 1988). Therefore, to induce cholesterol trafficking from the PM to ER, and hence the formation of MAM, we incubated hMNs^A4V^ and WT controls with SMase from *Bacillus cereus*, as described previously (Porn and Slotte, 1990; Slotte and Bierman, 1988; Slotte et al., 1990) (**Fig. S5**). As expected, SMase treatment induced the internalization of cholesterol (determined by increased filipin staining) and the formation and activation of MAM domains as measured by the synthesis and transfer of phospholipids (**Fig. S5 and S5B**), as well as by the increase in the expression of *MFN2* compared to non-treated cells (**Fig. 6A**). In support of a role for MAM in the regulation of pyruvate metabolism, treatment of hMNs^A4V^ with SMase rescued previously-observed defects in NADH-linked CI-OCR (**Fig. 6B**). This was consistent with the restoration of PDH activity, which promotes pyruvate oxidation (**Fig. 6C**), and a reduction in CPT1 activity, which would decrease FA import (**Fig. 6D**).

**Fig. 6.**
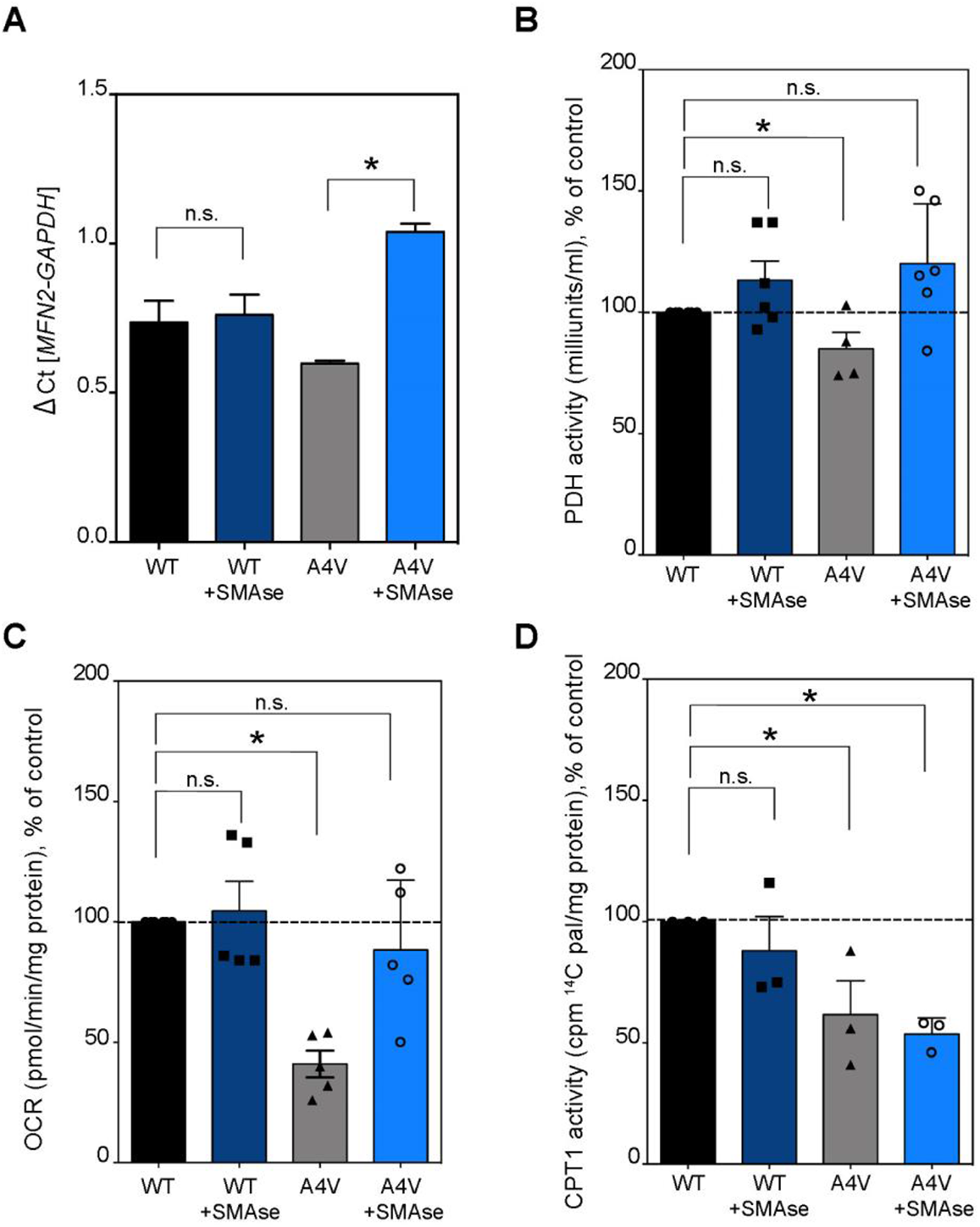
Stimulation of MAM formation reverses mitochondrial phenotypes in ALS motor neurons. Quantification of **(A)** *MFN2* mRNA expression, and **(B)** PDH activity in hMNs^A4V^ before and after sphingomyelinase (SMase) treatment compared to untreated WT cells (set at 100%). (Average ± SD of n=4-7 biological replicates using 2-4 technical replicates each; two-tailed t-test; α=0.05; *, p<0.05). (**C**) Quantification of OCR in hMNs^A4V^ using pyruvate as a mitochondrial substrate before and after treating with SMase relative to that in untreated WT cells (set at 100%) (Average ± SD of n=5-6 biological replicates using at least 3 technical replicates; two-tailed t-test; α=0.05; *, p<0.05). **(D)** CPT1 activity in hMNs^A4V^ before and after sphingomyelinase (SMase) treatment compared to untreated WT cells (set at 100%). (Average ± SD of n=3 biological replicates using 2-4 technical replicates each; two-tailed T-test; α=0.05; *, p<0.05).

Taken together, these results imply that MAM disruption in ALS precedes the alterations in glucose/pyruvate metabolism and the subsequent mitochondrial deficits.

## Discussion

We show here that, in the context of mutated SOD1 in ALS, the disruption of MAM impairs pyruvate metabolism and its use as mitochondrial fuel, which in turn induces a shift towards the use of fatty acids as an alternative substrate for ATP production. We posit that, when prolonged over time, this shift contributes to the mitochondrial respiratory defects characteristic of ALS. Therefore, our results lead us to conclude that the bioenergetic alterations observed in ALS-SOD1 mutants (and perhaps other forms of ALS as well) are downstream consequences of a prior impairment in the regulation of MAM. Of note, our results indicate that the spinal cord is affected earlier and more severely than is the brain, in agreement with the course of the disease. Although the reason behind these differences between SPC and brain is unknown, it is possible that the high energetic demands of motor neurons render SPC tissues more sensitive to the loss of metabolic flexibility caused by a progressive disruption in MAM domains, compared to other cell types and areas of the CNS.

MAM domains of the ER are cellular hubs for the regulation of several metabolic pathways in response to changes in the environment. Several reports have highlighted that, in addition to its role in lipid homeostasis and calcium regulation, MAM is an essential locus for the regulation of glucose metabolism and metabolic flexibility (Flis et al., 2018; Theurey and Rieusset, 2017; Townsend et al., 2020). Specifically, MAM integrity was shown to be essential for insulin and glucagon signaling via the recruitment and modulation of glycolytic enzymes such as hexokinase-II (Perry et al., 2020; Townsend et al., 2020). Moreover, the disruption of ER-mitochondria contacts and MAM domains was shown to be associated with impairments in pyruvate metabolism and insulin resistance (Rieusset, 2018), whereas increased MAM formation improved glucose tolerance (Tubbs et al., 2018a).

In support of previous reports (Sakai et al., 2021; Watanabe et al., 2016), our results show that ER-mitochondria communication and MAM functions are diminished in SOD1^G93A^ and SOD1^A4V^ models. We have also found that these models suffer from alterations in the lipid composition of their MAM domains as well as in reductions in the expression of the ER-mitochondria tether mitofusin-2. Remarkably, MAM defects have also been found in models of ALS carrying mutations in genes other than *SOD1*, such as *SIGMAR1*, *TARDP*, and *FUS* (De Vos et al., 2012; Parakh and Atkin, 2021; Sakai et al., 2021; Watanabe et al., 2016). In addition, stimulating the expression of known ER-mitochondria tethers, such as MFN2, was able to rescue mitochondrial abnormalities in fALS models with mutations in TDP-43 and SOD1 (Kim et al., 2016; Wang et al., 2013). These results argue that, at least in familial forms of the disease, disturbances in MAM could be a common and relevant alteration.

Based on our data and previous publications, we propose that in SOD1 mutants, defects in the regulation of MAM impair glucose metabolism and the production and oxidation of pyruvate, resulting in reductions in NADH-linked OxPhos. In turn, impairments in pyruvate utilization shift mitochondria towards the use of other carbon sources for ATP production, such as FAs. This shift is consistent with the mobilization of FAs from triglyceride stores and with alterations in glucose metabolism and insulin resistance that are characteristic of patients (Dorst et al., 2011; Mariosa et al., 2017; Pradat et al., 2010) and of mouse models of ALS (Palamiuc et al., 2015; Zhao et al., 2012). Moreover, the decreased use of pyruvate as a mitochondrial fuel source, and the reductions in LDH activity [lactate→pyruvate], could explain the high levels of lactate (and glucose) in the blood of ALS patients (Pradat et al., 2010; Siciliano et al., 2001). Furthermore, under conditions of low glucose utilization, excess glucose can be channeled through alternative, non-ATP-producing pathways, such as the formation of advanced glycation end products, synthesis of hexosamine, and astrocytic storage of glycogen, all of which are upregulated in ALS (Dodge et al., 2013; Juranek et al., 2015; Moll et al., 2020). Finally, a shift towards the use of FAs could also underlie the high levels of free FAs and triglyceride-rich lipoproteins found in blood and tissue samples from ALS patients (Area-Gomez et al., 2021; Dorst et al., 2011; Mariosa et al., 2017), and the observation that high-fat diets slow the progression of the disease (Mattson et al., 2007; Zhao et al., 2012).

This shift in substrate utilization can be detrimental for neuronal mitochondria. Specifically, the oxidation of FAs (and amino acids) results in higher levels of succinyl-CoA, succinate and FAD-based electron flux, as well as the activation of the electron-transferring flavoprotein (ETF), that ultimately converge increasing the levels of reduced CoQ (CoQH_2_) (Nicholls, 2013). In turn, elevations in the pool of (CoQH_2_) triggers RET and the reduction of NAD^+^ to NADH by CI (Chance and Hollunger, 1960), inducing higher NADH levels (and reduced NAD^+^:NADH ratios) (Kotlyar and Vinogradov, 1990). Moreover, these increases in succinate and stimulate complex CII, bypassing CI. As mentioned above, CII does not contribute to the proton-motive force, and thus results in lower ATP yields (Nicholls, 2013).

Our results confirm and extend these findings. Specifically, we have found that ALS-mutant cells and tissues present with lower ratios of NAD^+^:NADH, reduced activity of glycolytic enzymes, reduced OxPhos, increased ROS generation, and a shift in CI conformation to its dormant form – phenotypes that are all consistent with the persistent induction of RET. If true, this could help explain why cells from sporadic ALS and *C9ORF72* patients show higher membrane potential values compared to controls, despite displaying significant decreases in mitochondrial respiration (Kirk et al., 2014; Luengo et al., 2021; Onesto et al., 2016; Smith et al., 2019).

Under normal conditions, induction of RET allows the cell to adapt the structure and function of respiratory complexes to low-glucose conditions (Brand, 2016). However, when <u>prolonged over time</u>, RET can result in the loss of mitochondrial metabolic flexibility, bioenergetic defects, and oxidative damage, via increased ROS generation (Scialo et al., 2017; Stepanova et al., 2019a). Under RET, the reduction of NAD^+^ by CI results in the production of higher levels of NADH and superoxide (O_2_^-^), and subsequent elevations in *SOD1* mRNA expression and H_2_O_2_ production (Dell’Orco et al., 2016; Lambert et al., 2008; Pryde and Hirst, 2011); all of these phenotypes have been observed in ALS (Gagliardi et al., 2010). The elevations in H_2_O_2_, which can induce the expression of hypoxia-inducing factor 1α (HIF1α) (Kaewpila et al., 2008), could help explain the development of pseudo-hypoxic phenotypes (Selak et al., 2005) in the disease and the increased expression of HIF1α target genes frequently observed in ALS models (Sato et al., 2012, 2013).

It has also been shown that hypoxia causes the conversion of CI to the deactivate or “D form” (Chouchani et al., 2014; Gorenkova et al., 2013; Stepanova et al., 2019a). Although the “D form (CI-D)” can reversibly revert to the fully active “A form (CI-A)”, prolonged exposure to oxidative damage (e.g., ROS) can irreversibly inactivate CI and increase the dormant form (Drose et al., 2014; Drose et al., 2016). Notably, CI-D is more susceptible to covalent modifications such as nitrosylation and oxidation (Galkin and Moncada, 2007; Gorenkova et al., 2013), well-known contributors to ALS pathogenesis disease progression (Crow et al., 1997; Estevez et al., 1999).

In line with the idea that RET is activated in ALS tissues, our results show that CI activation is indeed delayed in SPC mitochondria from SOD1^G93A^ mice. This increase in the CI-D pool (due to a reduced transition from CI-D to CI-A; see **Fig. 4C**) results in higher rates of ROS production in the presence of pyruvate (Babot et al., 2014; Gorenkova et al., 2013), but *<u>not</u>* in the presence of succinate, as previous observed (Drose et al., 2016). This result could be explained mechanistically: whereas pyruvate generates NADH in the flavin site of CI-D whose excess electrons are shunted to produce ROS (**see Fig. 4D**), succinate-derived electrons (from CoQ) also arrive at CI-D but cannot proceed to the FMN- and therefore cannot generate ROS (Drose et al., 2016) (**see Fig 4, right panels**).

The progressive inactivation of CI due to prolonged FA overfeeding could also help explain the decreases in CI activity often reported during ALS (Ghiasi et al., 2012; Singh et al., 2021). One of the major findings of the current study is that the reduced D-to-A transition was detected <u>only</u> in SPC mitochondria. Brain, despite presenting increased FA oxidation, showed a higher pool of active CI in mutants compared to controls, but the mechanism(s) behind this phenotype is unknown. It is possible that the differential susceptibility of brain vs SPC is imposed by the different lipidic environment of each tissue, as shown in heart samples (Babot and Galkin, 2013; Loskovich et al., 2005).

Alternatively, the different metabolic phenotypes observed in brain and SPC under the same metabolic constraints can be the result of disparities in the expression or regulation of the components of the NAD^+^:NADH pathways and/or the shuttling of NAD^+^:NADH between mitochondria and the cytosol. In agreement, previous evidence has demonstrated that MNs are particularly sensitive to changes in NAD^+^:NADH ratio and that reductions in NAD^+^ could be a common phenomenon in motor neuron disorders (Lundt and Ding, 2021; Obrador et al., 2021). We also note that the potential build-up of NADH inside mitochondria could also induce a feedback inhibition of CPT1 and subsequent incomplete FA oxidation and increased production of short AC species in SPC tissues (Bartlett and Eaton, 2004).

Our findings lead us to propose the following model of ALS pathogenesis. In a healthy neuron, the formation of MAM domains favors pyruvate metabolism and NADH-fueled respiration via CI, with relatively <u>low</u> ROS generation (**Fig. 7**). In this scenario, the cell maintains a high NAD^+^:NADH ratio. In contrast, in ALS (**Fig. 7**), a disruption of MAM functionality induces a shift from pyruvate oxidation to FA β-oxidation, stimulating FADH_2_-CII respiration, and increasing the pool of reduced CoQH_2_. In this scenario, RET is triggered, resulting in a low NAD^+^:NADH ratio, high levels of ROS and changes in CI conformation (from CI-A → CI-D) (Drose et al., 2016) and function.

**Fig. 7.**
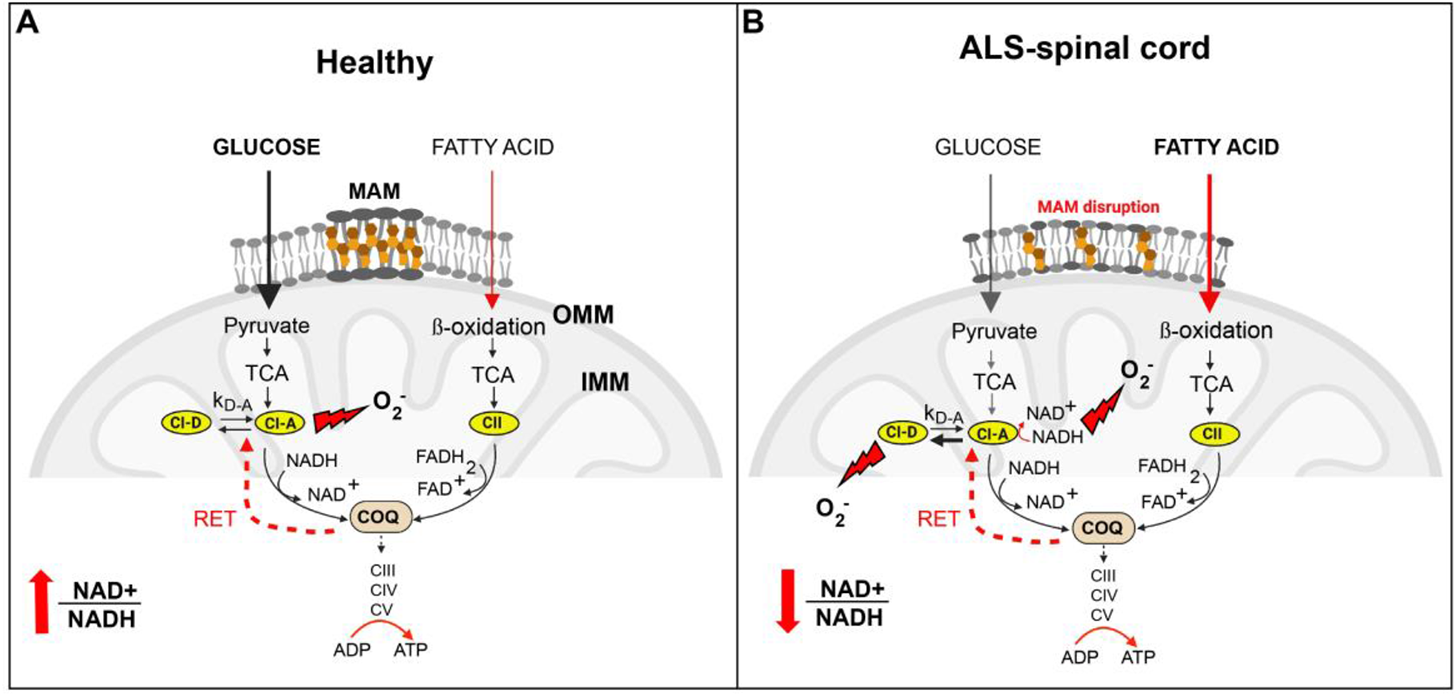
Model of mitochondrial metabolic dysfunction in ALS. MAM downregulation shifts mitochondrial metabolism in SOD1-mutant models and precedes mitochondrial defects in models of ALS. **(A)** In healthy neurons, mitochondria associate with the ER and this association promotes pyruvate metabolism, and thus NADH-OCR with relatively <u>low</u> ROS generation. Note that the cell maintains a high NAD^+^:NADH ratio to sustain the continuous production of NADH to fuel and ATP production. **(B)** In contrast, in the SPC of ALS mice, the disruption of MAM hinders pyruvate metabolism and induces a shift from pyruvate to other carbon sources such as FA, promoting increases in succinate and FADH_2_-OCR respiration and the transfer of electrons from CII to CoQ. Over time, this surge in FA oxidation increases the pool of reduced CoQ, triggering RET and the subsequent higher levels of ROS. In turn, the radicals generated in this process affects the kinetics of CI activation differentially in SPC as compared to brain (Fig.S6). In this scenario the SPC presents with a low NAD^+^:NADH ratio. The figure was created with BioRender (BioRender.com).

At the molecular level, we propose that CI activation (CI-D → CI-A transition) occurs faster in ALS brain than in controls but in SPC occurs more slowly than in controls (**Fig.S6**). This suggests that, in ALS, SPC CI is predominantly present in the D form, promoting ROS (**Fig. 7**).

Taken together, our data implicate disrupted ER-mitochondrial communication at the MAM as a primary event in the pathogenesis of ALS, and as a critical mediator of mitochondrial metabolic defects in this devastating disorder.

## Statements and Declarations

### Authors’ contributions

Conceived the project: EA-G and DL. Designed experiments: EA-G and DL. Provided with key reagents: JWS, ERL, MP and HW. Generated data for most of the experiments: DL, KAT, KK, KRV, JM and RRA. Critically assisted with complex I analysis: AG, AS and BY-S. Collected/analyzed lipidomics data: TDY and EA-G. Wrote the manuscript: DL and EA-G. Critically edited the manuscript: DL, RRA, AG and EA-G. Approved final version of the manuscript: all authors.

### Data availability

Source data files have been submitted to the journal as requested. The lipidomics datasets supporting the conclusions of this article are included as a supplemental file and have been submitted to appropriate data banks (www.metabolomicsworkbench.org). All reagents used in this work are commercially available, with manufacturer catalog numbers included in the Methods section.

### Ethics approval

All animal husbandry was conducted in accordance with the Guide for the Care and Use of Laboratory Animals published by the National Institutes of Health. Specific procedures were approved by the Institutional Animal Care and Use Committee at Columbia University protocol # AC-AABK5603.

## Acknowledgements

We thank Eric A. Schon and Serge Przedborski for critical discussions. We thank James Caicedo and Norma Romero for laboratory assistance. We thank Renu Nandakumar for assistance with the lipidomics analysis. Finally, we thank all members of the Motor Neuron Center at Columbia University Irving Medical Center for helpful discussions and support.

## Materials and Methods

### Animal and cell models

#### hSOD1 transgenic mice

All animal husbandry and experimentation were conducted in accordance with the National Institutes of Health Guide for Care and Use of Laboratory Animals. All procedures were approved by the Columbia University Institutional Animal Care and Use Committee (protocol # AC-AABK5603) and were consistent with the National Institutes of Health Guide for Care and Use of Laboratory Animals. Here we used B6SJLF1/J mice (SOD1*G93A)1Gur/J (002726) that have a high transgene copy number of the G93A-mutated hSOD1, and age-matched non-transgenics (NTg), from The Jackson Laboratories (Bar Harbor, ME). Genotypes were verified by PCR of tail DNA using the RedExtract-N-Amp Tissue PCR Kit (Sigma Aldrich 254-457-8).

G93A mice began exhibiting paralytic symptoms at 90 days of age and had mean lifespans of 120-130d. For biochemical and transcriptional assays, animals were sacrificed by decapitation at post-natal day 15 (P15), 30 (P30), 60 (P60), 90 (P90) or 120 (P120). For mitochondrial and enzymatic assays, brain and spinal cord (SPC) tissues were removed immediately and washed in cold phosphate-buffered saline (PBS) (HyClone SH30028.02) and homogenized or snap-frozen in liquid nitrogen or in dry ice for downstream analyses as described below. For histological studies, mice were anesthetized with 4% isoflurane gas and sacrificed via transcardial perfusion with PBS followed by 4% paraformaldehyde (PFA). For all studies, each experimental sample consisted of pooled tissues from at least two mice at a female/male ratio of 1:1. At least three technical replicates per group were assayed in every experiment, and every experiment was repeated at least 3 times for 3 biological replicates.

#### Human embryonic stem cell (hESC)-derived motor neurons (MNs)

We used the SOD1^+/A4V^ and SOD1^+/+^ isogenic cell lines described previously (Thams et al., 2019). Differentiation into human SPC MNs was also performed as described (Thams et al., 2019) with some modifications. The cell lines were maintained in mTeSR1 Plus Medium (Stemcell Technologies 100-0276 and 1 µM of Rock inhibitor (RI) (Tocris 1254). Differentiation was achieved by culturing the cells for 16 days in N2B27 media: a 1:1 ratio of Advanced DMEM/F12 medium (Life Technologies 12634028) containing N2 supplement (Life Technologies 17502048) and Neurobasal medium (Life Technologies 21103049), containing B27 supplement (without Vitamin A; Life Technologies 12587010), with 25 µM ß-mercaptoethanol and standard concentrations of pen/strep (Life Technologies 15140122) and L-glutamine (Life Technologies 25030081). Supplementation of N2B27 media with various factors was conducted as follows. **Day 0**: 10 ng/ml recombinant human FGF basic/FGF2/bFGF (146 aa) protein (R&D Systems 233-FB), 10 µM RI **(what is RI?)**, 20 µM SB431542 (Sigma S4217), 0.1 µM LDN-193189 (Stemgent 04-0074-02), 3 µM CHIR 99021 (Tocris 4423/10) and 10 µM ascorbic acid (AA) (Stemcell Technologies 72132). **Day 2**: 20 µM SB431542, 0.1 µM LDN-193189, 3 µM CHIR 99021, 10 µM AA, 100 nM retinoic acid (RA) (Tocris 1254) and 500 nM Smoothened Agonist (SAG Stemcell Technologies 73414). **Day 4**: 20 µM SB431542, 0.1 µM LDN-193189, 10 µM AA, 100 nM RA and 500 nM SAG. **Day 7**: 10 µM AA, 100 nM RA and 500 nM SAG. **Day 11**: 10 µM AA, 100 nM RA, 500 nM SAG, 10 ng/ml recombinant human brain-derived neurotrophic factor (BDNF) protein (R&D Systems 248-BDB-005), and 10 µM ɣ-secretase inhibitor (DAPT) (R&D Systems 2634/10). **Day 14**: 10 µM AA, 100 nM RA, 500 nM SAG, 10 ng/ml BDNF, and 10 µM DAPT. The cells were dissociated on **day 16** and plated on poly-L-lysine- and laminin-coated surfaces. The cells were differentiated *in vitro* for 2, 7 or 14 days in neuronal differentiation media [Neurobasal, N2, B27 with vitamin A (Life Technologies 17504 - 044), pen/strep, glutamine, non-essential amino acids (NEAA) (Life Technologies 11140076), antimitotic 1 µM UFdU, 10 mM uridine (Sigma-Aldrich U3750), 10 mM fluorodeoxyuridine (Sigma-Aldrich F0503), 10 ng/ml glial cell-derived neurotrophic factors (GDNF), 10 ng/ml BDNF, 10 ng/ml ciliary neurotrophic factor (CNTF), 25 µM glutamate (GluE), 25 µM ß-mercaptoethanol, 10 µM Forskolin, and 100 µM 3-isobutyl-1-methylxantine (IBMX)]. All cell lines used were routinely tested for mycoplasma following manufacturer instructions (Genlantis MY1100).

### Bioenergetics analyses

Mitochondrial fractions were purified from mouse tissue homogenates. Respirometry of isolated mitochondria from brain and SPC tissue and from cell cultures was performed using the Seahorse XFe24 Extracellular Flux Analyzer (Seahorse Bioscience; Agilent, Santa Clara, CA). Mitochondrial oxygen consumption rate (OCR) was determined as described previously (Agrawal et al., 2020; Pera et al., 2017). In summary, mice were sacrificed by decapitation at P15, P30, P60, P90 and P120. The brain and SPC were extracted immediately and washed in cold PBS. The tissues were homogenized in 10 volumes of homogenization buffer (210 mM mannitol, 70 mM sucrose, 5 mM HEPES and 1 mM EDTA) diluted 1:1 in Washing Buffer [210 mM mannitol, 70 mM sucrose, 5 mM HEPES and 1 mM EDTA, 0.5% FAF-BSA (Sigma #A7511) pH 7.2], and centrifuged at 900g for 10 min at 4°C. The remaining supernatant was centrifuged at 9000g for 10 min at 4°C. The resulting pellets were resuspended in homogenization buffer and centrifuged again at 9000 g for 10 min at 4°C and protein concentration was measured using the Quick Start Bradford Protein Assay Kit 1 (Bio-Rad 5000201). Mitochondrial respiration was measured in Mitochondrial Assay Solution (MAS, 70 mM sucrose, 220 mM mannitol, 5 mM KH_2_PO_4_, 5 mM MgCl_2_, 2 mM HEPES, 1 mM EGTA, 0.2% FAF-BSA, pH 7.4 adjusted with 1M KOH). Stock solutions containing 10X substrate stocks were prepared as follows: for Complex I-driven respiration, 50 mM pyruvate (Sigma-Aldrich P2256) + 50 mM malate (Sigma-Aldrich M8304); for Complex II-driven respiration, 50 mM succinate (Sigma S-2378) + 20 µM rotenone (Sigma-Aldrich 75351) (final concentration of each substrate was 5 mM for pyruvate/malate/succinate and 2 µM for rotenone). For complex I-mediated respiration, 8 μg protein was resuspended in MAS + CI substrates; for complex II-mediated respiration, 6 μg protein was resuspended in MAS + CII substrates in a final volume of 50 µL. The plate was centrifuged at 2000g for 5 min at 4°C, and 450 µL additional buffer was added to each well. In blank wells, 500 µL MAS was added. Oxygen consumption was measured at States 2, 3, 4, and 3-uncoupled (3U) after sequential addition of 3 mM ADP (Sigma-Aldrich A5285), 4 μM oligomycin (Sigma-Aldrich C75351), 6 μM carbonyl-cyanide p-triflouromethoxyphenylhydrazone (FCCP) (Sigma-Aldrich C2920) and 4.5 μM Antimycin A (Sigma-Aldrich A8674), respectively.

To measure OCR in whole cells, 50,000 cells were seeded in each well of a 24-well Seahorse assay plate, and differentiated to motor neurons for days *in vitro* (DIV) 2, DIV 7 and DIV14. OCR was measured under basal conditions (Seahorse media with 2 mM pyruvate and 25 mM glucose) and after the sequential addition of 1 μM oligomycin, 0.75 μM FCCP and 1 μM rotenone/1 μM antimycin A. Protein concentration was measured in each well after the experiment and all OCR values were normalized accordingly. All results were averages of at least 4 biological replicates each consisting of at least three technical replicates (assay wells) each.

#### Permeabilization assays

50,000 cells were seeded in each well of a 24-well Seahorse assay plate. The culture medium was replaced with MAS buffer containing 10 nM XF plasma membrane permeabilizer (PMP) reagent (Seahorse Bioscience 102504–100) with 5 mM malate + 5 mM pyruvate (for complex I assays) or 5 mM succinate + 2 μM rotenone (for complex II assays). OCR was measured first with no added substrates (state 2), and then after the sequential addition of 3 mm ADP (state 3), 4 μM oligomycin (state 4) and 6 μM FCCP (state 3-uncoupled). All results were averages of at least 3 biological replicates each consisting of at least three technical replicates (assay wells) each.

#### Co-staining for COX and SDH activities

Mice were sacrificed by decapitation at P60. The SPC was extracted and immediately snap-frozen in isopentane at −40°C for 35 sec. Following storage at −80°C, the SPCs were sectioned at 20 μm thickness using a CM3050S cryostat (Leica Biosystems, Buffalo Grove, IL). The sections were stored at −80°C until ready for sequential staining, at which point they were equilibrated at room temperature (RT) for 30 min before initiation of the staining protocol. The staining solution consisted of 2.7 mM 3,3-diaminobenzidine (DAB) (Sigma D56370), 0.09 mM cytochrome *c* from equine heart (Sigma C-7752), and 3U/ml catalase in 5 mM phosphate buffer, pH 7.4, filtered through Whatman paper. 100-200 µL was applied onto tissue sections for 40 min at 37°C in the dark, followed by 4 washes for 10 min each with ddH_2_O. Following completion of the COX staining protocol, the samples were subjected to SDH staining. The solution consisted of 5 mM EDTA, 1 mM KCN, 0.2 mM phenazine methosulfate (Sigma P9625), 50 mM succinic acid and 1.5 mM nitrotetrazolium blue chloride (Sigma N6876) (in 5 mM phosphate buffer, pH 7.6, filtered through Whatman paper). The stain was applied to the tissues for 40 min at 37°C in the dark, followed by 4 washes for 10 min each with ddH_2_O. The slides were coverslipped using warmed glycerin jelly as the mounting medium. Images were collected the next day using a Nikon Eclipse 80i brightfield microscope (Franco-Iborra and Tanji, 2020).

### Enzymatic activity assays

#### Complex II activity

This assay was conducted as previously described (Stepanova et al., 2019c). Complex II-dependent activity was measured in a plate reader (SpectraMax M2, Molecular Devices) recorded as the reduction of 2,6-dichloroindophenol (DCIP) as the chromophore (100 μM, 600 nm, ε = 21 mM^−1^ cm^−1^, blue to colorless) in HEPES buffer containing 60 µM DCIP, 40 µM decylubiquinone Q1, 0.5 µg/ml alamethicin (Cayman Chemical Company 11425), 2.5 µM rotenone and 20 mM succinate. When required, ubiquinol oxidation by the respiratory chain was inhibited by 1 mM KCN, and atpenin A5 was used to inhibit complex II. The rate of the corresponding blank was subtracted from each corresponding sample.

#### HK/PDH/LDH activities

Enzymatic activities were measured using commercial colorimetric quantification kits for hexokinase (HK; Sigma MAK091-1KT), pyruvate dehydrogenase (PDH; BioVision K679-100) and lactate dehydrogenase (LDH; BioVision K726-500). Briefly, 1X10^6^ or 10-20 mg of hMNs, 40-100 mg of fresh tissue total homogenates or 100-200 mg of crude mitochondrial fractions were isolated and resuspended in homogenization buffer following manufacturer instructions. Absorbance at 450 nm was measured using a Tecan Infinite 200 Pro plate reader at multiple time-points.

#### CPT1 activity

The carnitine palmitoyl transferase-1 (CPT1) assay was performed as described previously (Bremer, 1981) with modifications. Briefly, 200 mg of crude mitochondrial protein was added to the assay medium containing 20 mM HEPES (pH 7.3), 75 mM KCl, 2 mM KCN, 1% FAF-BSA, 70 mM palmitoyl-CoA and 0.25 mM L-[^3^H] carnitine, with or without 100 mM malonyl-CoA. Samples were incubated at 37°C for 10 min. The reaction was stopped by the addition of 0.5 ml of 1 M ice-cold perchloric acid. Mitochondria were centrifuged, the pellet was washed with 0.5 ml of 2 mM perchloric acid, and centrifugation was repeated. The resulting pellet was then resuspended in 0.8 mL ddH_2_O and extracted with 0.6 mL butanol. Three hundred microliters of the butanol phase were used for quantification of ^3^H radioactivity levels by liquid scintillation (Packard Tri-Carb 2900TR).

#### Sphingomyelinase treatment

Cells were incubated with 1 unit/ml of exogenous sphingomyelinase (SMase) from *Bacillus cereus* (Sigma S9396) for 1h at 37°C in neuronal differentiation media (described above). After incubation, the media was replaced and downstream analyses were performed.

#### Phospholipid synthesis and transfer

Cells were incubated for 2h in MEM (Gibco 11095098). The medium was replaced with MEM containing 2.5 μCi/mL ^3^H-serine and cell pellets were collected after 2, 4 or 6 hours. The cells were washed and collected in PBS, pelleted at 2500g for 5 min at 4°C and resuspended in 0.5 mL water, removing a small aliquot for protein quantification. Crude mitochondrial fractions were isolated using the same procedure as for the bioenergetics analyses. 50-100 µg of crude mitochondrial fractions were incubated in mitochondrial buffer containing 225 mM mannitol, 25mM HEPES and 1mM EGTA pH 7.4 supplemented with 5 μCi/mL ^3^H-serine and incubated at 37°C for 10 min. Lipid extraction was done using the Bligh and Dyer method (Bligh and Dyer, 1959). Briefly, 18 volumes of 1:2 chloroform/methanol were added to the samples and vortexed. After centrifugation at 8,000g for 5 min, the supernatant was transferred to a clean tube and 6 volumes of chilled chloroform followed by 5 volumes of chilled ddH_2_O were added to the tube, vortexed and centrifuged again at 9000 g for 2 min. The organic phase was transferred to a clean tube, concentrated, and re-extracted in 10 volumes of chilled chloroform, vortexed and centrifuged at 8000g for 2 min. Finally, the organic phase was transferred to a clean tube and dried under nitrogen. Dried lipids were resuspended in 100 μL 1:1 chloroform/methanol and spotted onto a silica thin layer chromatography (TLC) plate (catalog number). Phospholipids were separated using two solvents, composed of petroleum ether/diethyl ether/acetic acid 84:15:1 (v/v/v) and chloroform/methanol/acetic acid/water 60:50:1:4 (v/v/v/v). Development was performed by exposure of the plate to iodine vapor. The spots corresponding to the relevant phospholipids (identified using co-migrating standards) were scraped, exposed to liquid scintillation cocktail (catalog number), and counted in a scintillation counter (Packard Tri-Carb 2900TR).

#### Kinetics of complex-I D-form activation

Mice were sacrificed, and tissues (brains and SPC) were removed immediately, frozen in liquid nitrogen and stored at −80°C. Sixty to 80 mg of tissue was homogenized using a glass Dounce homogenizer in 2 ml of SET buffer (250 mM sucrose, 0.2 mM EDTA, 50 mM Tris-HCL), pH 8.8. The homogenization media was supplemented with 50 µg/ml of alamethicin and 1 mM ferricyanide (for preservation of physiological CI-A/CI-D ratios during isolation). Samples were centrifuged at 1,500g for 5 min at 40°C. The remaining supernatant was centrifuged at 20,000g for 20 min at 4°C and the resulting pellet was washed twice with SET buffer (pH 8.0) and subsequently resuspended in 100 ml of SET buffer (pH 7.5), aliquoted and stored at −80°C. At the time of the assay, mitochondrial samples were diluted to 2-5 mg/ml and resuspended in SET buffer (pH 8.5) supplemented with 5 mM MgCl_2_ and 15-30 μg alamethicin. The preparation was split into two samples: one incubated at 35°C for 30–60 minutes under constant shaking to obtain complex I in the D form, and a second one kept at 4°C. The activity was measured in SET buffer (pH 8.5) supplemented with 5 mM MgCl_2_, 15-30 μg alamethicin, and 5 μg cytochrome *c* containing 15–25 μg of mitochondrial protein/mL. The enzymatic activity of complex I in the D or A form was triggered by the addition of 140 μM NADH to the cuvette at 25°C and measured spectrophotometrically (Varian Cary 4000) as a decrease in absorption at 340 nm (ε340 nm = 6.22 mM^−1^ cm^−1^) (Galkin and Moncada, 2007; Stepanova et al., 2015). The first-rate order constant of the D-form to A-form transition was calculated in R version 3.4 (RStudio, PBC, Boston, MA) using semilogarithmic plots as described previously (Stepanova et al., 2015).

#### Measurement of mitochondrial H_2_O_2_ release in intact mitochondria

Mitochondrial respiration and H_2_O_2_ release were measured using a high-resolution respirometer (Oroboros Oxygraph-2k®) equipped with a two-channel fluorescence optical setup to use with the Amplex UltraRed and horseradish peroxidase assays (Stepanova and Galkin, 2020; Stepanova et al., 2019b). In brief, mitochondria (0.1 mg) were incubated in 2 ml of assay medium composed of 125 mM KCl, 0.2 mM EGTA, 20 mM HEPES-Tris, 4 mM KH_2_PO_4_ (pH 7.4), 2 mM MgCl_2_, 1 mg/ml BSA, 10 mM Amplex UltraRed, 10 U/ml SOD and 4 U/ml horseradish peroxidase at 37°C. To initiate respiration 2 mM malate and 5 mM pyruvate or 5 mM succinate and 1 mM glutamate were used. The H_2_O_2_ assays were calibrated by adding 100-200 pmol aliquots of fresh standard solution of H_2_O_2_ (ε_240nm_=46.3 M^-1^ cm^-1^) (Stepanova and Galkin, 2020; Stepanova et al., 2019b). ROS release was normalized by oxygen consumption in the sample at the same time of ROS release and the data were expressed as the average of at least 3 biological replicates each consisting of 3-4 technical replicates.

#### Quantification of NAD/NADH ratios

The NAD/NADH ratio was measured using the NAD/NADH Quantification Kit (Sigma-Aldrich MAK037) following manufacturer instructions. Briefly, to measure total NAD (NAD^+^ + NADH), 150 mg of cell homogenate or 20 mg of tissue were resuspended in 400 μL extraction buffer. After two cycles of freeze/thawing and vortexing, the cell extract was filtered using a 10-kDa cut-off spin filter and the filtrate was split into two 200-µL aliquots and kept on ice. To measure only NADH, 200 μL were incubated at 60°C for 30 min (to decompose NAD^+^), chilled and centrifuged at 9000g for 15 min. When tissues were assayed, 3 NTg and 3 G93A mice were sacrificed and their brains and SPCs were immediately removed and washed. Twenty mg of brain or SPC tissues were homogenized in 400 mL of extraction buffer and filtered using a 10-kDa filter. The filtrate was processed as described above for cell homogenates. Absorbance at 450 nm was measured using a Tecan Infinitive Pro 2000 plate reader at multiple time-points. The data were represented as the ratio of total NAD (NAD^+^ + NADH) to only NADH, and relative to controls.

#### Lipidomics analyses

Lipids were extracted from equal amounts of material (50-500 µg protein/sample). Lipid extracts were prepared via chloroform–methanol extraction, spiked with appropriate internal standards, and analyzed using a 6490 Triple Quadrupole LC/MS system (Agilent Technologies, Santa Clara, CA) as described previously (Chan et al., 2012). Lipids were separated by normal-phase HPLC using an Agilent Zorbax Rx-Sil column (inner diameter 2.1 × 100 mm) under the following conditions: mobile phase A (chloroform:methanol:1 M ammonium hydroxide, 89.9:10:0.1, v/v/v) and mobile phase B (chloroform:methanol:water:ammonium hydroxide, 55:39.9:5:0.1, v/v/v/v); 95% A for 2 min, linear gradient to 30% A over 18 min and held for 3 min, and linear gradient to 95% A over 2 min and held for 6 min. Quantification of lipid species was accomplished using multiple reaction monitoring (MRM) transitions that were developed in earlier studies (Chan et al., 2012) in conjunction with referencing of appropriate internal standards: ceramide d18:1/17:0 and sphingomyelin d18:1/12:0 (Avanti Polar Lipids, Alabaster, AL, USA). Values were represented as mole fraction with respect to total lipid (% molarity). For this, lipid mass (in moles) of any specific lipid was normalized by the total mass (in moles) of all the lipids measured (Chan et al., 2012). In addition, all of our results were further normalized by protein content.

#### Quantitative reverse transcription–polymerase chain reaction (qRT–PCR)

Total RNA was extracted from fresh tissues or cells using TRIzol Reagent (Invitrogen 15596-018) according to manufacturer instructions and was quantified by NanoDrop 2000 (Thermo Scientific). One to 5 µg of RNA were digested with DNase (Promega M6101) and reverse-transcribed to cDNA using the High-Capacity cDNA Reverse Transcription Kit (Applied Biosystems 4368813). Quantitative RT-PCR was performed in triplicate in a StepOnePlusTM Real-Time PCR System (Applied Biosystems 4376600). The expression of each gene under study was analyzed using specific predesigned TaqMan Probes (PGC-1α: *Ppargc1a* Mm01208835_m1; *PPARGC1A* Hs00173304_m1) (ThermoFisher 4331182) with *Gapdh/GAPDH* serving as the housekeeper gene (ThermoFisher 4352339E and 4352339E).

#### Subcellular fractionation and western blotting

Purification of MAM and mitochondria were performed and analyzed as described (Area-Gomez et al., 2009). Western blot analysis was performed as described (Larrea et al., 2019). For western blotting we used primary antibodies recognizing MFN2 (Abcam ab50838, Cambridge, United Kingdom), TOM20 (Santa Cruz sc-11415, Dallas, TX), VDAC1 (Abcam ab15895) and α-tubulin (Sigma-Aldrich T6199). To detect mitochondrial respiratory complexes, we used Total OXPHOS Rodent WB Antibody Cocktail (Abcam ab110413). As secondary antibodies, we used horseradish peroxidase-linked anti-rabbit IgG (Sigma-Aldrich NA934V) or anti-mouse IgG (Sigma-Aldrich NA931V).

### Fluorescent Microscopy

#### Filipin staining

hMNs were grown on coverslips and fixed with 4% PFA for 30 min. After 3 washes with 1X PBS, PFA was quenched by incubating the cells with 1.5 mg glycine/ml PBS for 10 min. The cells were stained with filipin complex (Sigma-Aldrich F9765) at a concentration of 0.5 mg/ml in 1X PBS with 10% FBS for 2h at room temperature. After extensive washes, coverslips were mounted with Fluoromount-G (ThermoFisher 00-4958-02) and imaged using a Zeiss Axiovert Fluorescence Microscope using a UV filter set [340–380 nm excitation, 40 nm dichroic, 430-nm long pass filter].

#### HB9-regulated GFP fluorescence

The hMNs^A4V^ were grown on coverslips and fixed with 4% PFA for 30 min at RT. The coverslips were mounted with Fluoromount-G and imaged using a Zeiss Axiovert Fluorescence Microscope using a UV filter set [488 nm excitation and 509 nm emission].

### Statistical analysis

All averages reflect three or more independent experiments. Each experiment consisted of different set of samples (control; NTg for mice, and WT for cells) that were measured at the same time within the same age group. Tests of significance employed Student’s t-test at an α=0.05 significance level (p<0.05), unless indicated otherwise. All error bars in figures represent SD among biological replicates. All statistical analysis were performed using GraphPad Prism (Version 6.0e) and each biological replicate was normalized by control values of the same replicate within the same time intervals.

## References

Acin-Perez, R., Carrascoso, I., Baixauli, F., Roche-Molina, M., Latorre-Pellicer, A., Fernandez-Silva, P., Mittelbrunn, M., Sanchez-Madrid, F., Perez-Martos, A., Lowell, C.A., et al. (2014). ROS-triggered phosphorylation of complex II by Fgr kinase regulates cellular adaptation to fuel use. Cell Metab 19, 1020–1033.

Adeva-Andany, M.M., Carneiro-Freire, N., Seco-Filgueira, M., Fernandez-Fernandez, C., and Mourino-Bayolo, D. (2019). Mitochondrial beta-oxidation of saturated fatty acids in humans. Mitochondrion 46, 73–90.

Agrawal, R.R., Tamucci, K.A., Pera, M., and Larrea, D. (2020). Assessing mitochondrial respiratory bioenergetics in whole cells and isolated organelles by microplate respirometry. Methods Cell Biol 155, 157–180.

Allen, S.P., Rajan, S., Duffy, L., Mortiboys, H., Higginbottom, A., Grierson, A.J., and Shaw, P.J. (2014). Superoxide dismutase 1 mutation in a cellular model of amyotrophic lateral sclerosis shifts energy generation from oxidative phosphorylation to glycolysis. Neurobiol Aging 35, 1499–1509.

Area-Gomez, E., de Groof, A.J., Boldogh, I., Bird, T.D., Gibson, G.E., Koehler, C.M., Yu, W.H., Duff, K.E., Yaffe, M.P., Pon, L.A., et al. (2009). Presenilins are enriched in endoplasmic reticulum membranes associated with mitochondria. Am J Pathol 175, 1810–1816.

Area-Gomez, E., Del Carmen Lara Castillo, M., Tambini, M.D., Guardia-Laguarta, C., de Groof, A.J., Madra, M., Ikenouchi, J., Umeda, M., Bird, T.D., Sturley, S.L., et al. (2012). Upregulated function of mitochondria-associated ER membranes in Alzheimer disease. EMBO J 31, 4106–4123.

Area-Gomez, E., Larrea, D., Yun, T., Xu, Y., Hupf, J., Zandkarimi, F., Chan, R.B., and Mitsumoto, H. (2021). Lipidomics study of plasma from patients suggest that ALS and PLS are part of a continuum of motor neuron disorders. Sci Rep 11, 13562.

Babot, M., Birch, A., Labarbuta, P., and Galkin, A. (2014). Characterisation of the active/de-active transition of mitochondrial complex I. Biochim Biophys Acta 1837, 1083–1092.

Babot, M., and Galkin, A. (2013). Molecular mechanism and physiological role of active-deactive transition of mitochondrial complex I. Biochem Soc Trans 41, 1325–1330.

Bartlett, K., and Eaton, S. (2004). Mitochondrial beta-oxidation. Eur J Biochem 271, 462–469.

Bartolome, F., Wu, H.C., Burchell, V.S., Preza, E., Wray, S., Mahoney, C.J., Fox, N.C., Calvo, A., Canosa, A., Moglia, C., et al. (2013). Pathogenic VCP mutations induce mitochondrial uncoupling and reduced ATP levels. Neuron 78, 57–64.

Belanger, M., Allaman, I., and Magistretti, P.J. (2011). Brain energy metabolism: focus on astrocyte-neuron metabolic cooperation. Cell Metab 14, 724–738.

Berthiaume, J.M., Kurdys, J.G., Muntean, D.M., and Rosca, M.G. (2019). Mitochondrial NAD(+)/NADH Redox State and Diabetic Cardiomyopathy. Antioxid Redox Signal 30, 375–398.

Bligh, E.G., and Dyer, W.J. (1959). A rapid method of total lipid extraction and purification. Can J Biochem Physiol 37, 911–917.

Bonilla, E., Sciacco, M., Tanji, K., Sparaco, M., Petruzzella, V., and Moraes, C.T. (1992). New morphological approaches to the study of mitochondrial encephalomyopathies. Brain Pathol 2, 113–119.

Brand, M.D. (2016). Mitochondrial generation of superoxide and hydrogen peroxide as the source of mitochondrial redox signaling. Free Radic Biol Med 100, 14–31.

Brand, M.D., and Nicholls, D.G. (2011). Assessing mitochondrial dysfunction in cells. Biochem J 435, 297–312.

Bremer, J. (1981). The effect of fasting on the activity of liver carnitine palmitoyltransferase and its inhibition by malonyl-CoA. Biochim Biophys Acta 665, 628–631.

Chan, R.B., Oliveira, T.G., Cortes, E.P., Honig, L.S., Duff, K.E., Small, S.A., Wenk, M.R., Shui, G., and Di Paolo, G. (2012). Comparative lipidomic analysis of mouse and human brain with Alzheimer disease. J Biol Chem 287, 2678–2688.

Chance, B., and Hollunger, G. (1960). Energy-linked reduction of mitochondrial pyridine nucleotide. Nature 185, 666–672.

Chouchani, E.T., Pell, V.R., Gaude, E., Aksentijevic, D., Sundier, S.Y., Robb, E.L., Logan, A., Nadtochiy, S.M., Ord, E.N.J., Smith, A.C., et al. (2014). Ischaemic accumulation of succinate controls reperfusion injury through mitochondrial ROS. Nature 515, 431–435.

Crow, J.P., Sampson, J.B., Zhuang, Y., Thompson, J.A., and Beckman, J.S. (1997). Decreased zinc affinity of amyotrophic lateral sclerosis-associated superoxide dismutase mutants leads to enhanced catalysis of tyrosine nitration by peroxynitrite. J Neurochem 69, 1936–1944.

Crugnola, V., Lamperti, C., Lucchini, V., Ronchi, D., Peverelli, L., Prelle, A., Sciacco, M., Bordoni, A., Fassone, E., Fortunato, F., et al. (2010). Mitochondrial respiratory chain dysfunction in muscle from patients with amyotrophic lateral sclerosis. Arch Neurol 67, 849–854.

de Brito, O.M., and Scorrano, L. (2008). Mitofusin 2: a mitochondria-shaping protein with signaling roles beyond fusion. Antioxid Redox Signal 10, 621–633.

De Vos, K.J., Morotz, G.M., Stoica, R., Tudor, E.L., Lau, K.F., Ackerley, S., Warley, A., Shaw, C.E., and Miller, C.C. (2012). VAPB interacts with the mitochondrial protein PTPIP51 to regulate calcium homeostasis. Hum Mol Genet 21, 1299–1311.

Dell’Orco, M., Milani, P., Arrigoni, L., Pansarasa, O., Sardone, V., Maffioli, E., Polveraccio, F., Bordoni, M., Diamanti, L., Ceroni, M., et al. (2016). Hydrogen peroxide-mediated induction of SOD1 gene transcription is independent from Nrf2 in a cellular model of neurodegeneration. Biochim Biophys Acta 1859, 315–323.

Divakaruni, A.S., Wallace, M., Buren, C., Martyniuk, K., Andreyev, A.Y., Li, E., Fields, J.A., Cordes, T., Reynolds, I.J., Bloodgood, B.L., et al. (2017). Inhibition of the mitochondrial pyruvate carrier protects from excitotoxic neuronal death. J Cell Biol 216, 1091–1105.

Dodge, J.C., Treleaven, C.M., Fidler, J.A., Tamsett, T.J., Bao, C., Searles, M., Taksir, T.V., Misra, K., Sidman, R.L., Cheng, S.H., et al. (2013). Metabolic signatures of amyotrophic lateral sclerosis reveal insights into disease pathogenesis. Proc Natl Acad Sci U S A 110, 10812–10817.

Dorst, J., Kuhnlein, P., Hendrich, C., Kassubek, J., Sperfeld, A.D., and Ludolph, A.C. (2011). Patients with elevated triglyceride and cholesterol serum levels have a prolonged survival in amyotrophic lateral sclerosis. J Neurol 258, 613–617.

Drose, S., Brandt, U., and Wittig, I. (2014). Mitochondrial respiratory chain complexes as sources and targets of thiol-based redox-regulation. Biochim Biophys Acta 1844, 1344–1354.

Drose, S., Stepanova, A., and Galkin, A. (2016). Ischemic A/D transition of mitochondrial complex I and its role in ROS generation. Biochim Biophys Acta 1857, 946–957.

Estevez, A.G., Crow, J.P., Sampson, J.B., Reiter, C., Zhuang, Y., Richardson, G.J., Tarpey, M.M., Barbeito, L., and Beckman, J.S. (1999). Induction of nitric oxide-dependent apoptosis in motor neurons by zinc-deficient superoxide dismutase. Science 286, 2498–2500.

Flis, D.J., Dzik, K., Kaczor, J.J., Halon-Golabek, M., Antosiewicz, J., Wieckowski, M.R., and Ziolkowski, W. (2018). Swim Training Modulates Skeletal Muscle Energy Metabolism, Oxidative Stress, and Mitochondrial Cholesterol Content in Amyotrophic Lateral Sclerosis Mice. Oxid Med Cell Longev 2018, 5940748.

Franco-Iborra, S., and Tanji, K. (2020). Histochemical and immunohistochemical staining methods to visualize mitochondrial proteins and activity. Methods Cell Biol 155, 247–270.

Gagliardi, S., Cova, E., Davin, A., Guareschi, S., Abel, K., Alvisi, E., Laforenza, U., Ghidoni, R., Cashman, J.R., Ceroni, M., et al. (2010). SOD1 mRNA expression in sporadic amyotrophic lateral sclerosis. Neurobiol Dis 39, 198–203.

Galkin, A., and Moncada, S. (2007). S-nitrosation of mitochondrial complex I depends on its structural conformation. J Biol Chem 282, 37448–37453.

Ghiasi, P., Hosseinkhani, S., Noori, A., Nafissi, S., and Khajeh, K. (2012). Mitochondrial complex I deficiency and ATP/ADP ratio in lymphocytes of amyotrophic lateral sclerosis patients. Neurol Res 34, 297–303.

Gorenkova, N., Robinson, E., Grieve, D.J., and Galkin, A. (2013). Conformational change of mitochondrial complex I increases ROS sensitivity during ischemia. Antioxid Redox Signal 19, 1459–1468.

Guaras, A., Perales-Clemente, E., Calvo, E., Acin-Perez, R., Loureiro-Lopez, M., Pujol, C., Martinez-Carrascoso, I., Nunez, E., Garcia-Marques, F., Rodriguez-Hernandez, M.A., et al. (2016). The CoQH2/CoQ Ratio Serves as a Sensor of Respiratory Chain Efficiency. Cell Rep 15, 197–209.

Guardia-Laguarta, C., Area-Gomez, E., Rub, C., Liu, Y., Magrane, J., Becker, D., Voos, W., Schon, E.A., and Przedborski, S. (2014). alpha-Synuclein is localized to mitochondria-associated ER membranes. J Neurosci 34, 249–259.

Hall, E.D., Oostveen, J.A., and Gurney, M.E. (1998). Relationship of microglial and astrocytic activation to disease onset and progression in a transgenic model of familial ALS. Glia 23, 249–256.

Hasselbalch, S.G., Knudsen, G.M., Jakobsen, J., Hageman, L.P., Holm, S., and Paulson, O.B. (1994). Brain metabolism during short-term starvation in humans. J Cereb Blood Flow Metab 14, 125–131.

Hirst, J. (2013). Mitochondrial complex I. Annu Rev Biochem 82, 551–575.

Juranek, J.K., Daffu, G.K., Wojtkiewicz, J., Lacomis, D., Kofler, J., and Schmidt, A.M. (2015). Receptor for Advanced Glycation End Products and its Inflammatory Ligands are Upregulated in Amyotrophic Lateral Sclerosis. Front Cell Neurosci 9, 485.

Kaewpila, S., Venkataraman, S., Buettner, G.R., and Oberley, L.W. (2008). Manganese superoxide dismutase modulates hypoxia-inducible factor-1 alpha induction via superoxide. Cancer Res 68, 2781–2788.

Kawamata, H., Peixoto, P., Konrad, C., Palomo, G., Bredvik, K., Gerges, M., Valsecchi, F., Petrucelli, L., Ravits, J.M., Starkov, A., et al. (2017). Mutant TDP-43 does not impair mitochondrial bioenergetics in vitro and in vivo. Mol Neurodegener 12, 37.

Kim, J.Y., Jang, A., Reddy, R., Yoon, W.H., and Jankowsky, J.L. (2016). Neuronal overexpression of human VAPB slows motor impairment and neuromuscular denervation in a mouse model of ALS. Hum Mol Genet 25, 4661–4673.

Kirk, K., Gennings, C., Hupf, J.C., Tadesse, S., D’Aurelio, M., Kawamata, H., Valsecchi, F., Mitsumoto, H., Groups, A.P.C.S., and Manfredi, G. (2014). Bioenergetic markers in skin fibroblasts of sporadic amyotrophic lateral sclerosis and progressive lateral sclerosis patients. Ann Neurol 76, 620–624.

Kotlyar, A.B., and Vinogradov, A.D. (1990). Slow active/inactive transition of the mitochondrial NADH-ubiquinone reductase. Biochim Biophys Acta 1019, 151–158.

Koves, T.R., Ussher, J.R., Noland, R.C., Slentz, D., Mosedale, M., Ilkayeva, O., Bain, J., Stevens, R., Dyck, J.R., Newgard, C.B., et al. (2008). Mitochondrial overload and incomplete fatty acid oxidation contribute to skeletal muscle insulin resistance. Cell Metab 7, 45–56.

Lambert, A.J., Buckingham, J.A., and Brand, M.D. (2008). Dissociation of superoxide production by mitochondrial complex I from NAD(P)H redox state. FEBS Lett 582, 1711–1714.

Larrea, D., Pera, M., Gonnelli, A., Quintana-Cabrera, R., Akman, H.O., Guardia-Laguarta, C., Velasco, K.R., Area-Gomez, E., Dal Bello, F., De Stefani, D., et al. (2019). MFN2 mutations in Charcot-Marie-Tooth disease alter mitochondria-associated ER membrane function but do not impair bioenergetics. Hum Mol Genet 28, 1782–1800.

Lingwood, D., and Simons, K. (2010). Lipid rafts as a membrane-organizing principle. Science 327, 46–50.

Loskovich, M.V., Grivennikova, V.G., Cecchini, G., and Vinogradov, A.D. (2005). Inhibitory effect of palmitate on the mitochondrial NADH:ubiquinone oxidoreductase (complex I) as related to the active-de-active enzyme transition. Biochem J 387, 677–683.

Ludtmann, M.H.R., Arber, C., Bartolome, F., de Vicente, M., Preza, E., Carro, E., Houlden, H., Gandhi, S., Wray, S., and Abramov, A.Y. (2017). Mutations in valosin-containing protein (VCP) decrease ADP/ATP translocation across the mitochondrial membrane and impair energy metabolism in human neurons. J Biol Chem 292, 8907–8917.

Luengo, A., Li, Z., Gui, D.Y., Sullivan, L.B., Zagorulya, M., Do, B.T., Ferreira, R., Naamati, A., Ali, A., Lewis, C.A., et al. (2021). Increased demand for NAD(+) relative to ATP drives aerobic glycolysis. Mol Cell 81, 691–707 e696.

Lundt, S., and Ding, S. (2021). NAD(+) Metabolism and Diseases with Motor Dysfunction. Genes (Basel) 12.

Marangi, G., and Traynor, B.J. (2015). Genetic causes of amyotrophic lateral sclerosis: new genetic analysis methodologies entailing new opportunities and challenges. Brain Res 1607, 75–93.

Mariosa, D., Hammar, N., Malmstrom, H., Ingre, C., Jungner, I., Ye, W., Fang, F., and Walldius, G. (2017). Blood biomarkers of carbohydrate, lipid, and apolipoprotein metabolisms and risk of amyotrophic lateral sclerosis: A more than 20-year follow-up of the Swedish AMORIS cohort. Ann Neurol 81, 718–728.

Martinez-Reyes, I., and Chandel, N.S. (2020). Mitochondrial TCA cycle metabolites control physiology and disease. Nat Commun 11, 102.

Mattiazzi, M., D’Aurelio, M., Gajewski, C.D., Martushova, K., Kiaei, M., Beal, M.F., and Manfredi, G. (2002). Mutated human SOD1 causes dysfunction of oxidative phosphorylation in mitochondria of transgenic mice. J Biol Chem 277, 29626–29633.

Mattson, M.P., Cutler, R.G., and Camandola, S. (2007). Energy intake and amyotrophic lateral sclerosis. Neuromolecular Med 9, 17–20.

McGarry, J.D., and Foster, D.W. (1974). The metabolism of (minus)-octanoylcarnitine in perfused livers from fed and fasted rats. Evidence for a possible regulatory role of carnitine acyltransferase in the control of ketogenesis. J Biol Chem 249, 7984–7990.

Mergenthaler, P., Lindauer, U., Dienel, G.A., and Meisel, A. (2013). Sugar for the brain: the role of glucose in physiological and pathological brain function. Trends Neurosci 36, 587–597.

Moll, T., Shaw, P.J., and Cooper-Knock, J. (2020). Disrupted glycosylation of lipids and proteins is a cause of neurodegeneration. Brain 143, 1332–1340.

Montesinos, J., Area-Gomez, E., and Schlame, M. (2020). Analysis of phospholipid synthesis in mitochondria. Methods Cell Biol 155, 321–335.

Naon, D., Zaninello, M., Giacomello, M., Varanita, T., Grespi, F., Lakshminaranayan, S., Serafini, A., Semenzato, M., Herkenne, S., Hernandez-Alvarez, M.I., et al. (2016). Critical reappraisal confirms that Mitofusin 2 is an endoplasmic reticulum-mitochondria tether. Proc Natl Acad Sci U S A 113, 11249–11254.

Nicholls, D.G.a.F., S.T. (2013). Bioenergetics4. Elsevier

Obrador, E., Salvador, R., Marchio, P., Lopez-Blanch, R., Jihad-Jebbar, A., Rivera, P., Valles, S.L., Banacloche, S., Alcacer, J., Colomer, N., et al. (2021). Nicotinamide Riboside and Pterostilbene Cooperatively Delay Motor Neuron Failure in ALS SOD1(G93A) Mice. Mol Neurobiol 58, 1345–1371.

Onesto, E., Colombrita, C., Gumina, V., Borghi, M.O., Dusi, S., Doretti, A., Fagiolari, G., Invernizzi, F., Moggio, M., Tiranti, V., et al. (2016). Gene-specific mitochondria dysfunctions in human TARDBP and C9ORF72 fibroblasts. Acta Neuropathol Commun 4, 47.

Owen, O.E., Morgan, A.P., Kemp, H.G., Sullivan, J.M., Herrera, M.G., and Cahill, G.F., Jr. (1967). Brain metabolism during fasting. J Clin Invest 46, 1589–1595.

Palamiuc, L., Schlagowski, A., Ngo, S.T., Vernay, A., Dirrig-Grosch, S., Henriques, A., Boutillier, A.L., Zoll, J., Echaniz-Laguna, A., Loeffler, J.P., et al. (2015). A metabolic switch toward lipid use in glycolytic muscle is an early pathologic event in a mouse model of amyotrophic lateral sclerosis. EMBO Mol Med 7, 526–546.

Panov, A., Kubalik, N., Zinchenko, N., Hemendinger, R., Dikalov, S., and Bonkovsky, H.L. (2011). Respiration and ROS production in brain and spinal cord mitochondria of transgenic rats with mutant G93a Cu/Zn-superoxide dismutase gene. Neurobiol Dis 44, 53–62.

Parakh, S., and Atkin, J.D. (2021). The Mitochondrial-associated ER membrane (MAM) compartment and its dysregulation in Amyotrophic Lateral Sclerosis (ALS). Semin Cell Dev Biol 112, 105–113.

Pera, M., Larrea, D., Guardia-Laguarta, C., Montesinos, J., Velasco, K.R., Agrawal, R.R., Xu, Y., Chan, R.B., Di Paolo, G., Mehler, M.F., et al. (2017). Increased localization of APP-C99 in mitochondria-associated ER membranes causes mitochondrial dysfunction in Alzheimer disease. EMBO J 36, 3356–3371.

Perry, R.J., Zhang, D., Guerra, M.T., Brill, A.L., Goedeke, L., Nasiri, A.R., Rabin-Court, A., Wang, Y., Peng, L., Dufour, S., et al. (2020). Glucagon stimulates gluconeogenesis by INSP3R1-mediated hepatic lipolysis. Nature 579, 279–283.

Porn, M.I., and Slotte, J.P. (1990). Reversible effects of sphingomyelin degradation on cholesterol distribution and metabolism in fibroblasts and transformed neuroblastoma cells. Biochem J 271, 121–126.

Pradat, P.F., Bruneteau, G., Gordon, P.H., Dupuis, L., Bonnefont-Rousselot, D., Simon, D., Salachas, F., Corcia, P., Frochot, V., Lacorte, J.M., et al. (2010). Impaired glucose tolerance in patients with amyotrophic lateral sclerosis. Amyotroph Lateral Scler 11, 166–171.

Pryde, K.R., and Hirst, J. (2011). Superoxide is produced by the reduced flavin in mitochondrial complex I: a single, unified mechanism that applies during both forward and reverse electron transfer. J Biol Chem 286, 18056–18065.

Randle, P.J., Garland, P.B., Hales, C.N., and Newsholme, E.A. (1963). The glucose fatty-acid cycle. Its role in insulin sensitivity and the metabolic disturbances of diabetes mellitus. Lancet 1, 785–789.

Reyes, E.T., Perurena, O.H., Festoff, B.W., Jorgensen, R., and Moore, W.V. (1984). Insulin resistance in amyotrophic lateral sclerosis. J Neurol Sci 63, 317–324.

Rieusset, J. (2018). The role of endoplasmic reticulum-mitochondria contact sites in the control of glucose homeostasis: an update. Cell Death Dis 9, 388.

Robb, E.L., Hall, A.R., Prime, T.A., Eaton, S., Szibor, M., Viscomi, C., James, A.M., and Murphy, M.P. (2018). Control of mitochondrial superoxide production by reverse electron transport at complex I. J Biol Chem 293, 9869–9879.

Robberecht, W., and Philips, T. (2013). The changing scene of amyotrophic lateral sclerosis. Nat Rev Neurosci 14, 248–264.

Rowland, L.P., and Shneider, N.A. (2001). Amyotrophic lateral sclerosis. N Engl J Med 344, 1688–1700.

Sakai, S., Watanabe, S., Komine, O., Sobue, A., and Yamanaka, K. (2021). Novel reporters of mitochondria-associated membranes (MAM), MAMtrackers, demonstrate MAM disruption as a common pathological feature in amyotrophic lateral sclerosis. FASEB J 35, e21688.

Sato, K., Morimoto, N., Kurata, T., Mimoto, T., Miyazaki, K., Ikeda, Y., and Abe, K. (2012). Impaired response of hypoxic sensor protein HIF-1alpha and its downstream proteins in the spinal motor neurons of ALS model mice. Brain Res 1473, 55–62.

Sato, K., Morimoto, N., Kurata, T., Mimoto, T., Miyazaki, K., Ikeda, Y., and Abe, K. (2013). Impaired hypoxic sensor Siah-1, PHD3, and FIH system in spinal motor neurons of an amyotrophic lateral sclerosis mouse model. J Neurosci Res 91, 285–291.

Scaricamazza, S., Salvatori, I., Giacovazzo, G., Loeffler, J.P., Rene, F., Rosina, M., Quessada, C., Proietti, D., Heil, C., Rossi, S., et al. (2020). Skeletal-Muscle Metabolic Reprogramming in ALS-SOD1(G93A) Mice Predates Disease Onset and Is A Promising Therapeutic Target. iScience 23, 101087.

Schonfeld, P., and Reiser, G. (2013). Why does brain metabolism not favor burning of fatty acids to provide energy? Reflections on disadvantages of the use of free fatty acids as fuel for brain. J Cereb Blood Flow Metab 33, 1493–1499.

Schooneman, M.G., Vaz, F.M., Houten, S.M., and Soeters, M.R. (2013). Acylcarnitines: reflecting or inflicting insulin resistance? Diabetes 62, 1–8.

Scialo, F., Fernandez-Ayala, D.J., and Sanz, A. (2017). Role of Mitochondrial Reverse Electron Transport in ROS Signaling: Potential Roles in Health and Disease. Front Physiol 8, 428.

Selak, M.A., Armour, S.M., MacKenzie, E.D., Boulahbel, H., Watson, D.G., Mansfield, K.D., Pan, Y., Simon, M.C., Thompson, C.B., and Gottlieb, E. (2005). Succinate links TCA cycle dysfunction to oncogenesis by inhibiting HIF-alpha prolyl hydroxylase. Cancer Cell 7, 77–85.

Siciliano, G., Pastorini, E., Pasquali, L., Manca, M.L., Iudice, A., and Murri, L. (2001). Impaired oxidative metabolism in exercising muscle from ALS patients. J Neurol Sci 191, 61–65.

Singh, T., Jiao, Y., Ferrando, L.M., Yablonska, S., Li, F., Horoszko, E.C., Lacomis, D., Friedlander, R.M., and Carlisle, D.L. (2021). Neuronal mitochondrial dysfunction in sporadic amyotrophic lateral sclerosis is developmentally regulated. Sci Rep 11, 18916.

Slotte, J.P., and Bierman, E.L. (1988). Depletion of plasma-membrane sphingomyelin rapidly alters the distribution of cholesterol between plasma membranes and intracellular cholesterol pools in cultured fibroblasts. Biochem J 250, 653–658.

Slotte, J.P., Tenhunen, J., and Porn, I. (1990). Effects of sphingomyelin degradation on cholesterol mobilization and efflux to high-density lipoproteins in cultured fibroblasts. Biochim Biophys Acta 1025, 152–156.

Smith, E.F., Shaw, P.J., and De Vos, K.J. (2019). The role of mitochondria in amyotrophic lateral sclerosis. Neurosci Lett 710, 132933.

Speijer, D. (2011). Oxygen radicals shaping evolution: why fatty acid catabolism leads to peroxisomes while neurons do without it: FADH(2)/NADH flux ratios determining mitochondrial radical formation were crucial for the eukaryotic invention of peroxisomes and catabolic tissue differentiation. Bioessays 33, 88–94.

Speijer, D., Manjeri, G.R., and Szklarczyk, R. (2014). How to deal with oxygen radicals stemming from mitochondrial fatty acid oxidation. Philos Trans R Soc Lond B Biol Sci 369, 20130446.

Stanley, W.C., Recchia, F.A., and Lopaschuk, G.D. (2005). Myocardial substrate metabolism in the normal and failing heart. Physiol Rev 85, 1093–1129.

Stepanova, A., and Galkin, A. (2020). Measurement of mitochondrial H2O2 production under varying O2 tensions. Methods Cell Biol 155, 273–293.

Stepanova, A., Konrad, C., Guerrero-Castillo, S., Manfredi, G., Vannucci, S., Arnold, S., and Galkin, A. (2019a). Deactivation of mitochondrial complex I after hypoxia-ischemia in the immature brain. J Cereb Blood Flow Metab 39, 1790–1802.

Stepanova, A., Konrad, C., Manfredi, G., Springett, R., Ten, V., and Galkin, A. (2019b). The dependence of brain mitochondria reactive oxygen species production on oxygen level is linear, except when inhibited by antimycin A. J Neurochem 148, 731–745.

Stepanova, A., Sosunov, S., Niatsetskaya, Z., Konrad, C., Starkov, A.A., Manfredi, G., Wittig, I., Ten, V., and Galkin, A. (2019c). Redox-Dependent Loss of Flavin by Mitochondrial Complex I in Brain Ischemia/Reperfusion Injury. Antioxid Redox Signal 31, 608–622.

Stepanova, A., Valls, A., and Galkin, A. (2015). Effect of monovalent cations on the kinetics of hypoxic conformational change of mitochondrial complex I. Biochim Biophys Acta 1847, 1085–1092.

Steyn, F.J., Li, R., Kirk, S.E., Tefera, T.W., Xie, T.Y., Tracey, T.J., Kelk, D., Wimberger, E., Garton, F.C., Roberts, L., et al. (2020). Altered skeletal muscle glucose-fatty acid flux in amyotrophic lateral sclerosis. Brain Commun 2, fcaa154.

Stoica, R., De Vos, K.J., Paillusson, S., Mueller, S., Sancho, R.M., Lau, K.F., Vizcay-Barrena, G., Lin, W.L., Xu, Y.F., Lewis, J., et al. (2014). ER-mitochondria associations are regulated by the VAPB-PTPIP51 interaction and are disrupted by ALS/FTD-associated TDP-43. Nat Commun 5, 3996.

Szelechowski, M., Amoedo, N., Obre, E., Leger, C., Allard, L., Bonneu, M., Claverol, S., Lacombe, D., Oliet, S., Chevallier, S., et al. (2018). Metabolic Reprogramming in Amyotrophic Lateral Sclerosis. Sci Rep 8, 3953.

Tabassum, N.K., I.S.; Ibn Asaduzzaman, S.A.; Maniha, S.M.; Fayz, A.H.; Zakaria, A.; Noor, R. (2020). A Review on the Possible Leakage of Electrons through the Electron Transport Chain within Mitochondria. J Biomed Res Environ Sci. 1, 105–113.

Tefera, T.W., Steyn, F.J., Ngo, S.T., and Borges, K. (2021). CNS glucose metabolism in Amyotrophic Lateral Sclerosis: a therapeutic target? Cell Biosci 11, 14.

Thams, S., Lowry, E.R., Larraufie, M.H., Spiller, K.J., Li, H., Williams, D.J., Hoang, P., Jiang, E., Williams, L.A., Sandoe, J., et al. (2019). A Stem Cell-Based Screening Platform Identifies Compounds that Desensitize Motor Neurons to Endoplasmic Reticulum Stress. Mol Ther 27, 87–101.

Theurey, P., and Rieusset, J. (2017). Mitochondria-Associated Membranes Response to Nutrient Availability and Role in Metabolic Diseases. Trends Endocrinol Metab 28, 32–45.

Townsend, L.K., Brunetta, H.S., and Mori, M.A.S. (2020). Mitochondria-associated ER membranes in glucose homeostasis and insulin resistance. Am J Physiol Endocrinol Metab 319, E1053–E1060.

Tubbs, E., Axelsson, A.S., Vial, G., Wollheim, C.B., Rieusset, J., and Rosengren, A.H. (2018a). Sulforaphane improves disrupted ER-mitochondria interactions and suppresses exaggerated hepatic glucose production. Mol Cell Endocrinol 461, 205–214.

Tubbs, E., Chanon, S., Robert, M., Bendridi, N., Bidaux, G., Chauvin, M.A., Ji-Cao, J., Durand, C., Gauvrit-Ramette, D., Vidal, H., et al. (2018b). Disruption of Mitochondria-Associated Endoplasmic Reticulum Membrane (MAM) Integrity Contributes to Muscle Insulin Resistance in Mice and Humans. Diabetes 67, 636–650.

Vance, J.E. (1990). Phospholipid synthesis in a membrane fraction associated with mitochondria. J Biol Chem 265, 7248–7256.

Vance, J.E. (2014). MAM (mitochondria-associated membranes) in mammalian cells: lipids and beyond. Biochim Biophys Acta 1841, 595–609.

Wang, W., Li, L., Lin, W.L., Dickson, D.W., Petrucelli, L., Zhang, T., and Wang, X. (2013). The ALS disease-associated mutant TDP-43 impairs mitochondrial dynamics and function in motor neurons. Hum Mol Genet 22, 4706–4719.

Watanabe, S., Ilieva, H., Tamada, H., Nomura, H., Komine, O., Endo, F., Jin, S., Mancias, P., Kiyama, H., and Yamanaka, K. (2016). Mitochondria-associated membrane collapse is a common pathomechanism in SIGMAR1- and SOD1-linked ALS. EMBO Mol Med 8, 1421–1437.

Wiedemann, F.R., Manfredi, G., Mawrin, C., Beal, M.F., and Schon, E.A. (2002). Mitochondrial DNA and respiratory chain function in spinal cords of ALS patients. J Neurochem 80, 616–625.

Worth, A.J., Basu, S.S., Snyder, N.W., Mesaros, C., and Blair, I.A. (2014). Inhibition of neuronal cell mitochondrial complex I with rotenone increases lipid beta-oxidation, supporting acetyl-coenzyme A levels. J Biol Chem 289, 26895–26903.

Zhao, W., Varghese, M., Vempati, P., Dzhun, A., Cheng, A., Wang, J., Lange, D., Bilski, A., Faravelli, I., and Pasinetti, G.M. (2012). Caprylic triglyceride as a novel therapeutic approach to effectively improve the performance and attenuate the symptoms due to the motor neuron loss in ALS disease. PLoS One 7, e49191.

