## Supplementary Figures and Table 1 for "Altered MAM function shifts mitochondrial metabolism in SOD1-mutant models of ALS": DL. Larrea et al, 2022. Supplementary Figures V1.docx

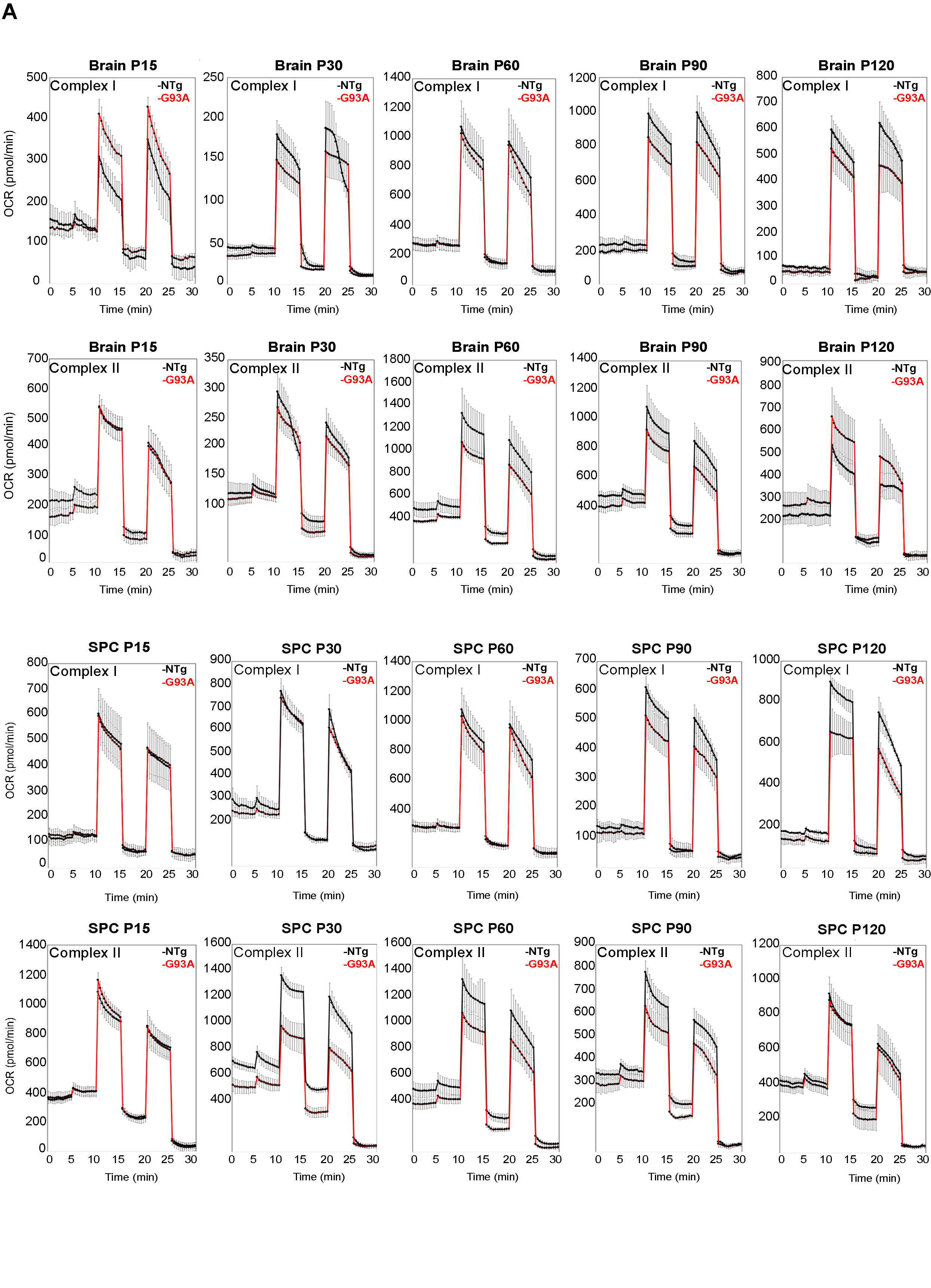
**Supplementary Figures**

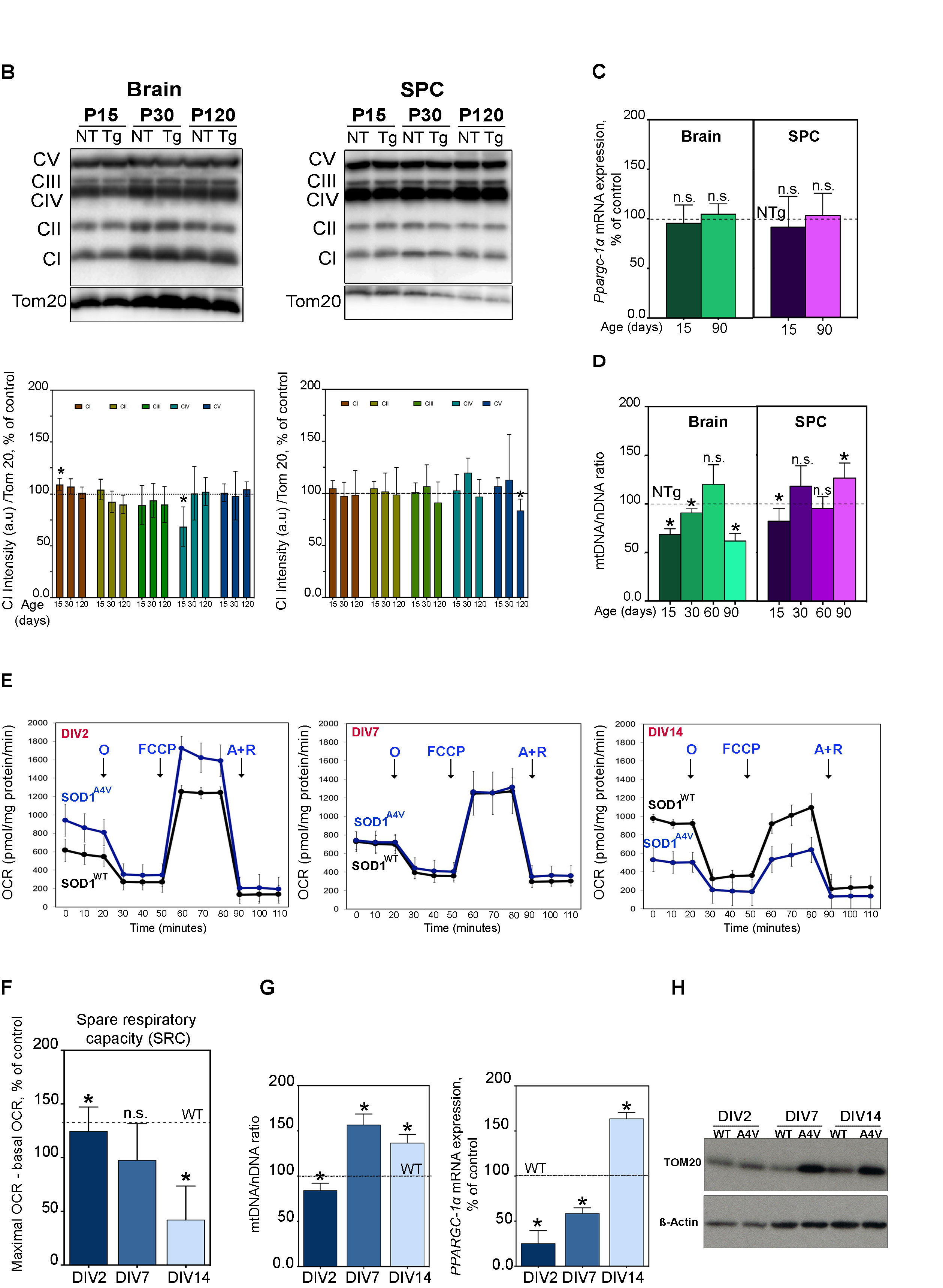

**Supplementary Figure 1. (A)** Representative Seahorse plots of OCR point by point at the indicated stages (P) in mitochondria isolated from brain and SPC of SOD1^G93A^ mice (red traces) and NTg controls (black traces) at P15, P30, P60, P90 and P120. Initial respiration in the presence of added substrates (i.e., pyruvate+malate to measure CI-mediated respiration or succinate+rotenone to measure CII-mediated respiration) was measured after the addition of oligomycin (Oligo); FCCP and antimycin. All experiments were performed with at least 3 replicates; *p < 0.05. **(B)** Western blot of representative subunits from respiratory complexes I (NDUFB8, ~20 kDa), II (SDHB, ~30 kDa), III (UQCRC2, ~48 kDa), IV (MTCO2, ~40 kDa), and V (ATP5A, ~55 kDa) in SOD1^G93A^ brain and SPC relative to NTg controls. Quantification of the western blots is shown at right, normalized to TOM20 protein expression and reported relative to NTg control levels. **(C)** Expression of *Ppargc1a*) (encoding Pgc1-α) mRNA in brain and SPC from SOD1^G93A^ mice normalized to *Gapdh* mRNA and reported relative to the NTg control average (set at 100%, dotted line), at the indicated ages, in days. Data are shown as the average ± SD (two-tailed t-test; ⍺=0.05; *, p<0.05). **(D)** Quantitation of mtDNA/nuclear DNA ratio in SOD1^G93A^ mice relative to the NTg control average (set at 100%, dotted line). The results show the average of n=3 biological replicates. Data are shown as the average ± SD; (two-tailed t-test; ⍺=0.05; *, p<0.05). **(E)** Representative Seahorse plots of OCR in hMNs^A4V^ and WT whole cells after the addition of oligomycin, FCCP, and antimycin + rotenone, at the indicated DIVs. **(F)** Spare respiratory capacity calculated as the maximal respiration (FCCP) minus basal respiration in mutant cells compared to WT (set as 100% of control; dotted line). The results show the average of n=4-5 biological replicates and each experiment was performed using at least 3 technical replicates. Data are shown as the average ± SD; (two-tailed t-test; ⍺=0.05; *, p<0.05). **(G)** Quantitation of mtDNA/nuclear DNA ratio relative to control average (left panel) and expression of total *PPARGC1A* mRNA (right panel) in hMNs^A4V^ relative to WT at the indicated DIVs. **(H)** representative Western blot of hMNs proteins to detect TOM20, normalized to b-actin. The results show the average of n=3-4 biological replicates and each experiment was performed using at least 3 technical replicates. Data are shown as the average ± SD; (two-tailed t-test; ⍺=0.05; *, p<0.05)

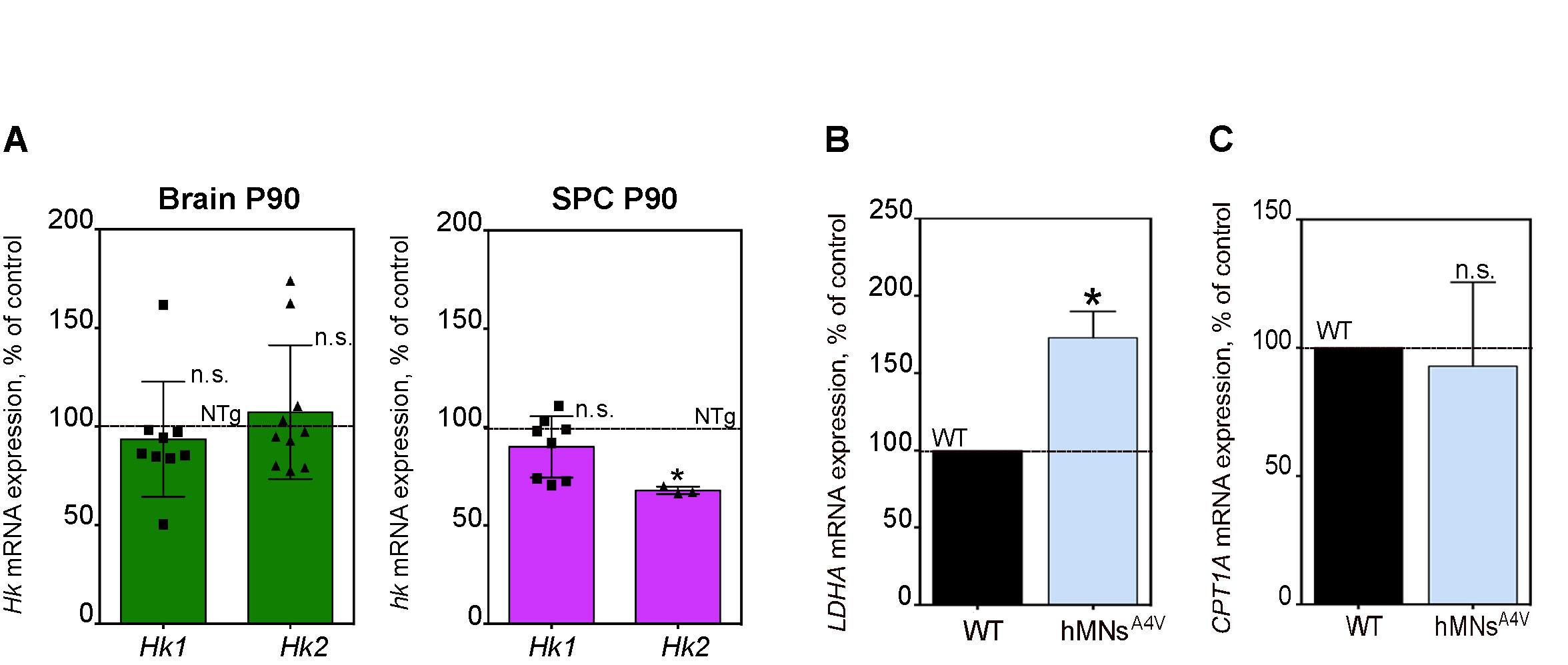
**Supplementary Fig. 2. Real time PCR. (A)** Quantification of *Hk1* and *Hk2* mRNA expression in brain and SPC from SOD1^G93A^ mice normalized to *Gapdh*. **(B-C)** Quantification of *LDHA* and *CPT1A* mRNA expression in hMNs^A4V^ normalized to *GAPDH*. All assays are averages of 3-4 biological replicates each consisting of 3 technical replicates. Data are shown as mean ± SD. Statistical comparisons are made to NTg (for mouse experiments) or WT (for hMN experiments) controls (two-tailed t-test; ⍺=0.05; *, p<0.05).

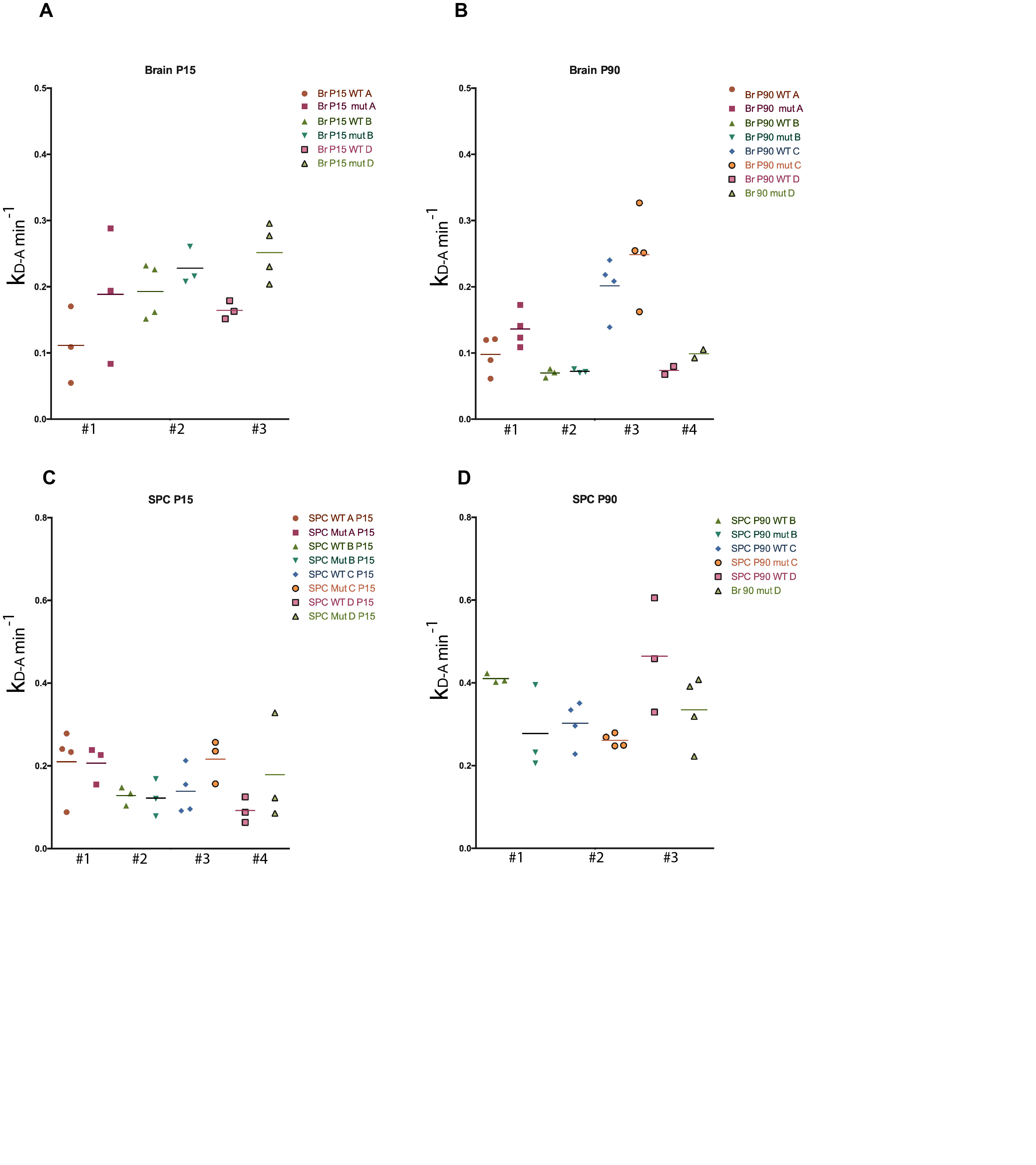

**Supplementary Fig. 3. Defects in NADH levels and complex I activation in mitochondria from SOD1^G93A^ mice.** **(A-D)** Analysis of the D-to-A transition constant for CI, expressed as min^-1^, as k_D-A_ min^-1^. Shown are individual experiments and technical replicates in brain and SPC at P15 and P90 as indicated in color code at the right of each set of experiments.

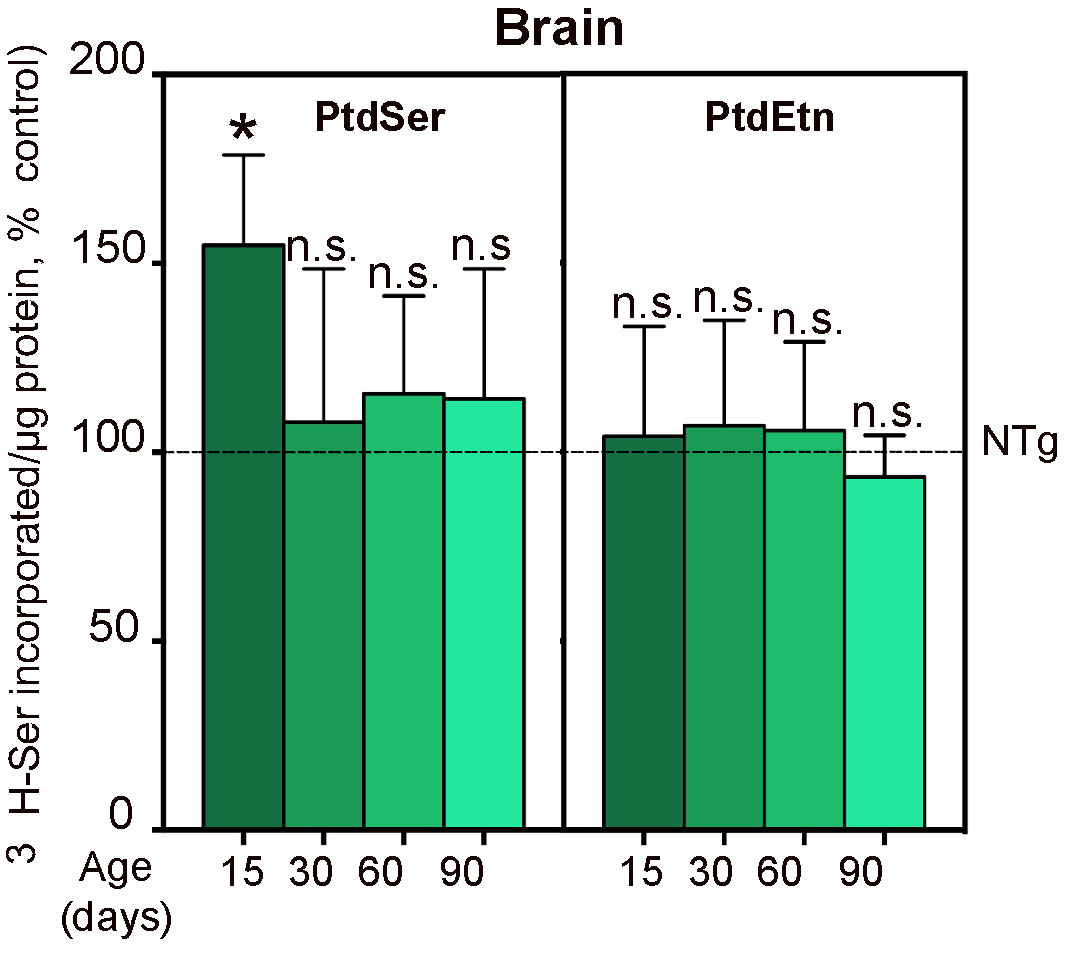

**Supplementary Fig. 4. MAM activity in SOD1^G93A^ brain.** Quantification of MAM activity in brain tissues isolated from SOD1^G93A^ mice at the indicated ages, in days. MAM activity was measured by the synthesis and transfer of phospholipids between the ER and mitochondria, or the incorporation after 20 min of ^3^H-Ser into ^3^H-PtdSer and ^3^H-PtdEtn, relative to NTg controls (set at 100%; dotted line). All assays are averages of 3-4 biological replicates, each consisting of 3 technical replicates. Data are shown as mean ± SD. Statistical comparisons are made to NTg (for mouse experiments) or WT (for hMN experiments) controls (two-tailed t-test; ⍺=0.05; *, p<0.05).

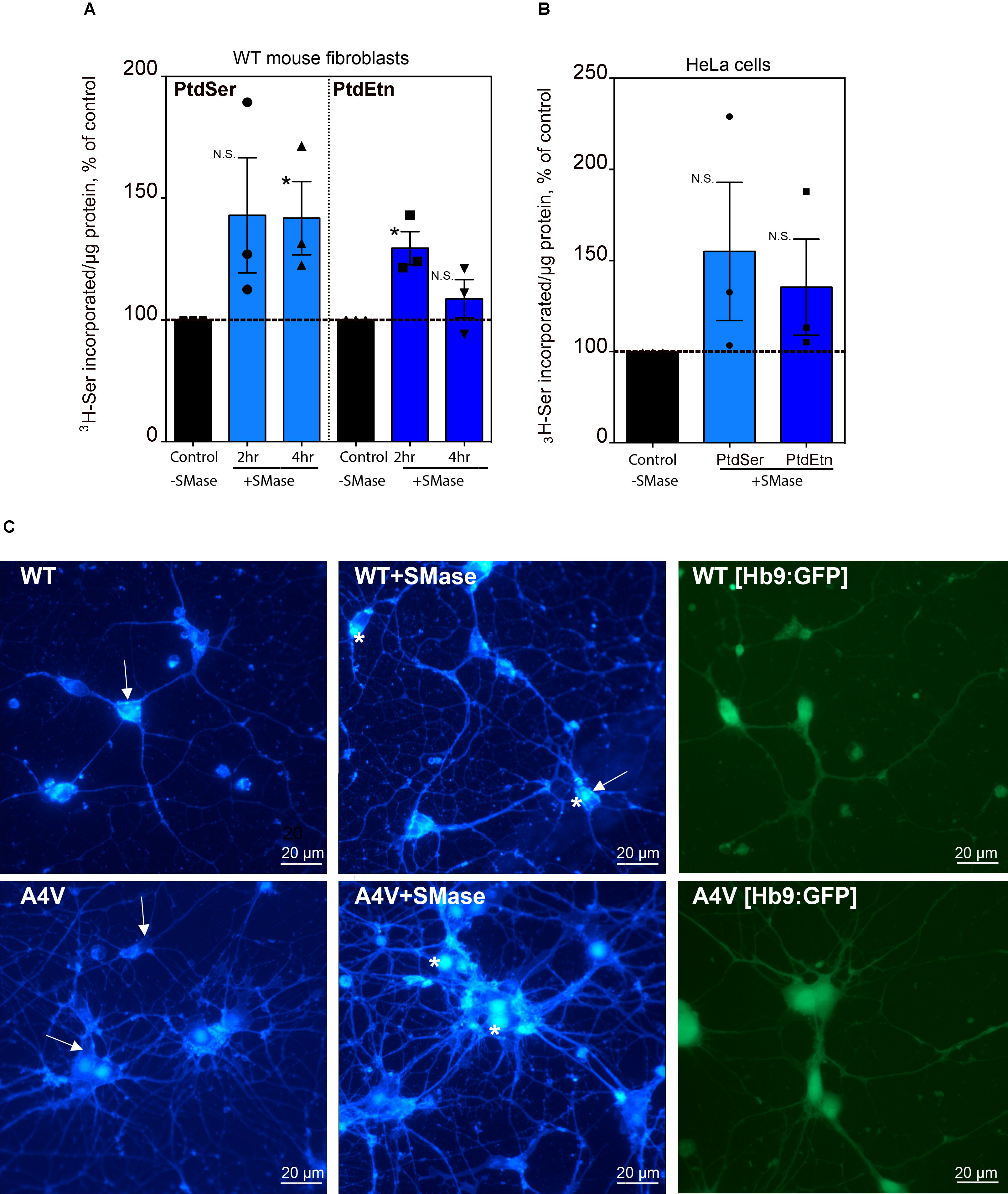

**Supplementary Fig. 5. Induction of MAM upon treatment with exogenous sphingomyelinase.** Quantification of MAM activity by the synthesis and transfer of phospholipids between ER and mitochondria in **(A)** mouse fibroblasts at 2h and 4 h and **(B)** HeLa cells at 4 h relative to non-treated cells (dotted lines, set at 100%). (Average ± SD of n = 3 biological replicates; two-tailed t-test; ⍺=0.05; *, p<0.05). **(C)** Confocal images of cholesterol in hMNs^A4V^ and WT controls. Cells were treated with and without SMase and stained with filipin (left and middle columns, as indicated). Arrows indicate presence or absence of filipin staining in the membrane, whereas asterisks indicate the accumulation cholesterol. The right column shows representative pictures of both cell lines expressing GFP under the control of a specific motor neuron promoter, HB9.

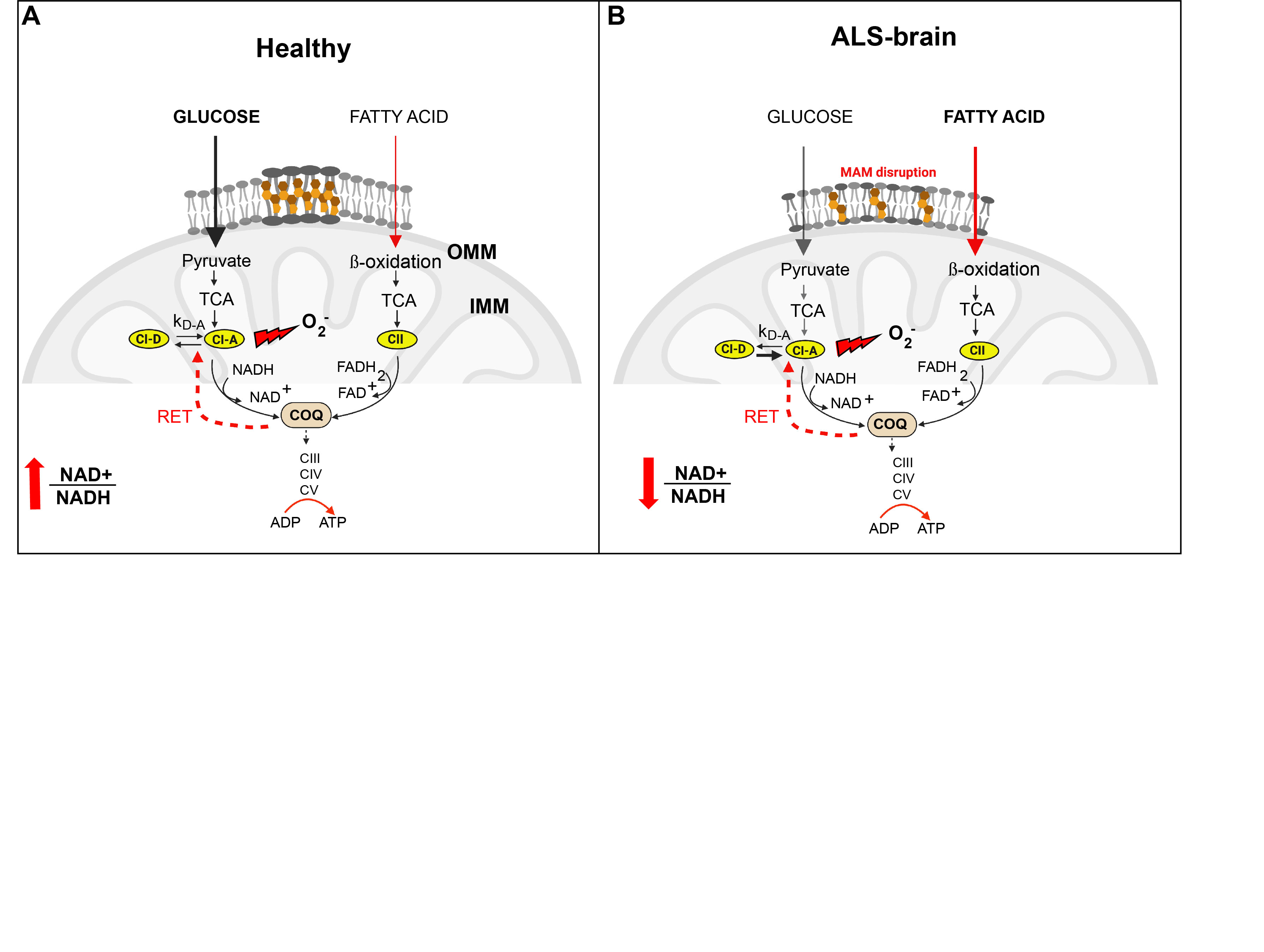

**Supplementary Fig. 6. Continuation of the model for MAM downregulation in the brain of ALS mouse models.** As stated in Fig. 7, in healthy neurons, the formation of MAM domains promotes pyruvate metabolism, and thus NADH-OCR respiration with the generation of low ROS levels. In contrast, in the context of SOD1 mutations, the disruption of MAM hinders pyruvate metabolism and induces a shift from pyruvate to other carbon sources such as FA, promoting increases in succinate and FADH_2_-OCR respiration and the transfer of electrons from CII to CoQ. Over time, this surge in FA consumption increases the pool of reduced CoQ, triggerring RET and the subsequent higher levels of ROS. In turn, the radicals generated in this process affects the kinetics of CI activation in SPC (**Fig. 7).** In this scenario the cell presents with low NAD^+^:NADH ratio. Conversely, in brain tissues even under RET conditions and lower NAD^+^:NADH ratios, ALS-brain mitochondria are able to maintain a pool of active CI in forward mode (i.e., transferring electrons from NADH to CoQ), generating low levels of ROS associated with the oxidation of NADH-linked substrates (Drose et al., 2016).

**Supplementary Table 1:** OCR absolute values for respirometry experiments (pmol/min/mg mitochondrial protein)

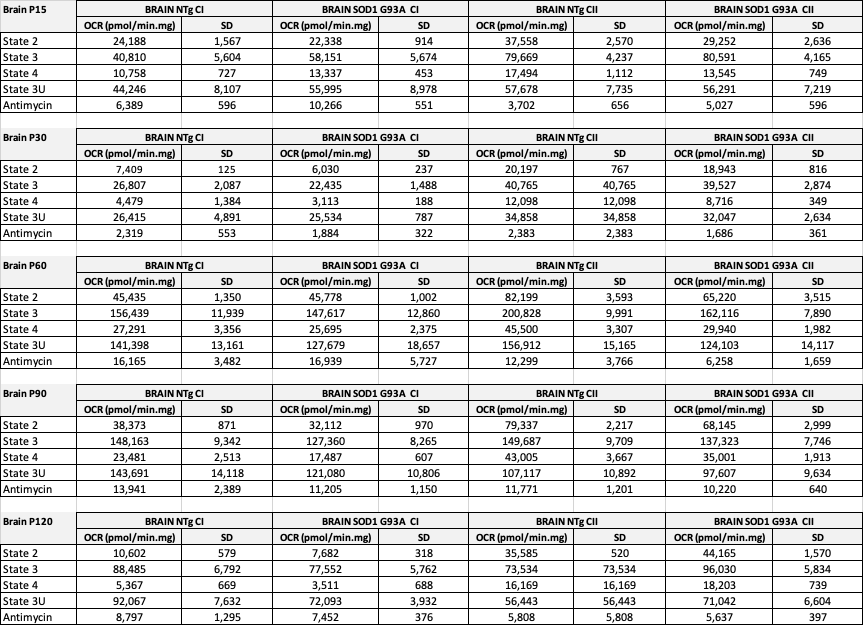

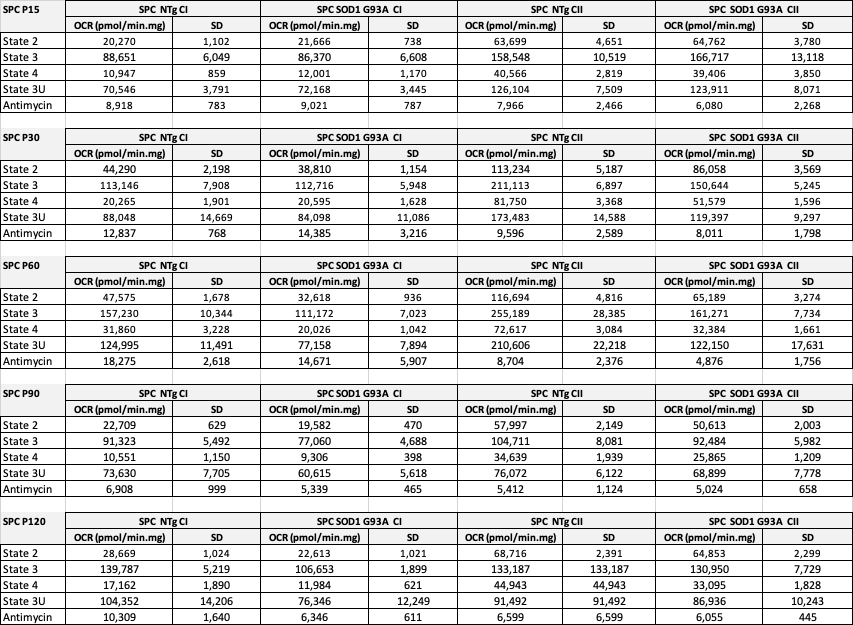
